# A framework for Frizzled-G protein coupling and implications to the Wnt-PCP signaling pathways

**DOI:** 10.1101/2023.07.08.548223

**Authors:** Zhibin Zhang, Xi Lin, Ling Wei, Yiran Wu, Lu Xu, Lijie Wu, Xiaohu Wei, Arthur Wang, Suwen Zhao, Xiangjia Zhu, Fei Xu

## Abstract

The ten Frizzled receptors (FZDs) are essential in Wnt signaling and play important roles in embryonic development and tumorigenesis. Among these, FZD6 is closely associated with lens development. Understanding FZD activation mechanism is key to unlock these emerging targets. Here we present the cryo-EM structures of FZD6 and FZD3 which are known to relay non-canonical Wnt-PCP (planar cell polarity) signaling pathways as well as FZD1 in their G protein-coupled (active) and G protein-free (inactive) states, respectively. Comparison of the three inactive/active pairs unveiled a shared activation framework among all ten FZDs. Mutagenesis along with imaging and functional analysis on the human lens epithelial tissues suggested potential crosstalk between G-protein binding and Wnt-PCP signaling pathways. Together, this study provides an integrated understanding of FZD structure and function, and lays the foundation for developing therapeutic modulators to activate or inhibit FZD signaling for a range of disorders including cancers and cataracts.

## Introduction

Frizzled proteins (FZDs) are primary Wnt receptors belonging to the class F G protein-coupled receptors (GPCRs) that consist of ten FZDs and one Smoothened (SMO)^1, 2^. 19 mammalian Wnts can bind to ten FZDs thereby activating different downstream pathways and regulating key events in embryonic development as well as tissue homeostasis and regeneration in the adult organism^3^. Therefore, dysregulation of these receptors can lead to various human diseases ranging from cancer, developmental defects to metabolic and neurological disorders^4, 5^. All FZDs contain a highly conserved cysteine-rich domain (CRD), the orthosteric binding site for Wnt ligands, at the extracellular region^6^. The CRD domain is followed by a flexible linker varying in length in different FZDs and a compact hinge domain before the transmembrane domain^7^ (Extended Data Fig. 1a). Unlike most GPCRs, FZD signaling was initially divided into two major pathways, being either dependent (canonical signaling) or independent (non-canonical signaling) on the accumulation of the transcription regulator β-catenin. The two non-canonical Wnt signaling pathways are the planar cell polarity (PCP) pathway and the Ca^2+^ pathway^8^.

**Fig. 1.**
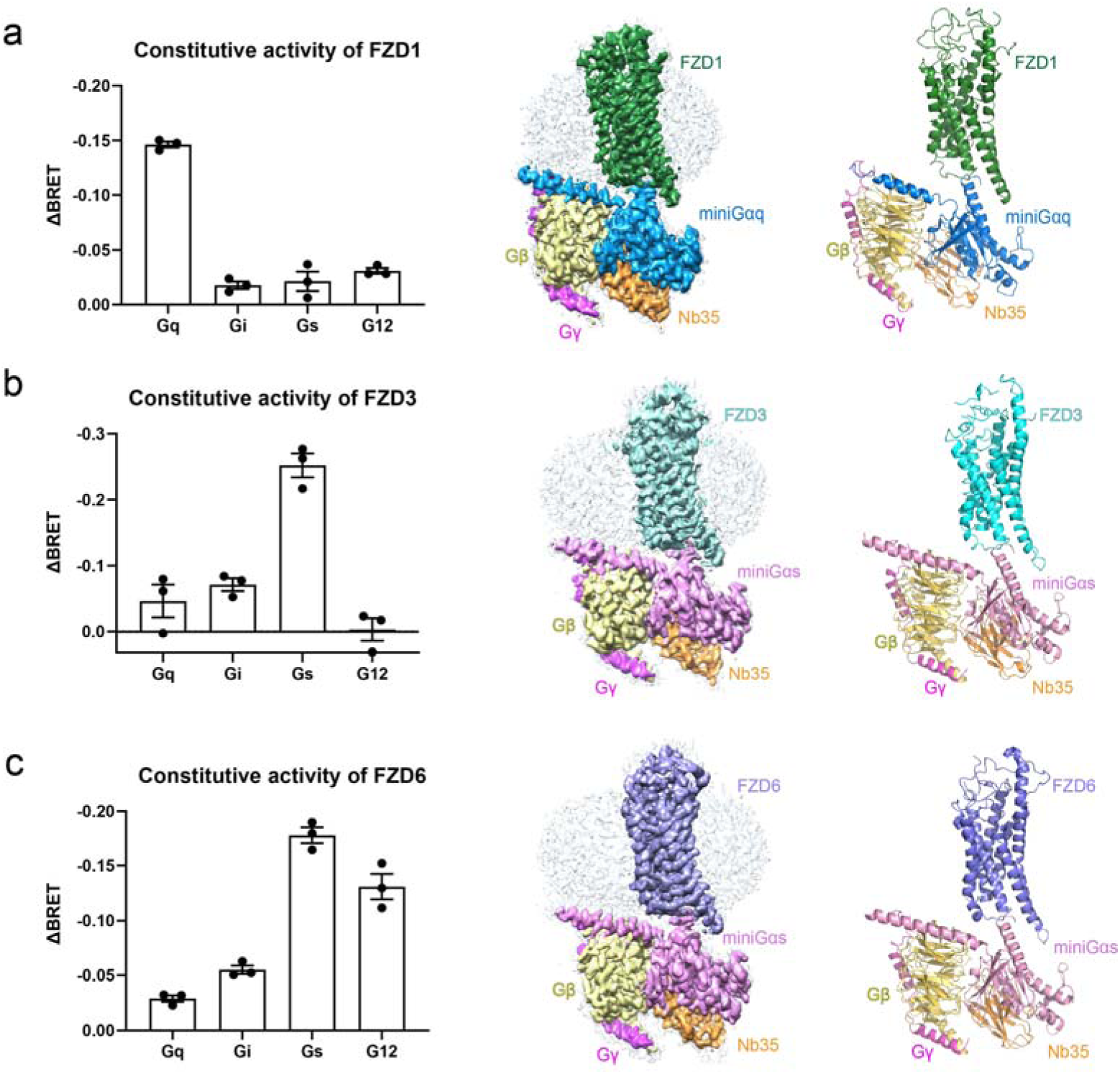
| Cryo-EM structures of the FZD1-Gq, FZD3-Gs and FZD6-Gs complexes. **a-c**, The BRET-based Gq, Gi, Gs and G12 biosensor is used to examine the constitutive activity of FZD1, FZD3 and FZD6 (left). Cryo-EM map (middle) and cartoon representation of the atomic model (right) of the FZD1-Gq (**a**), FZD3-Gs (**b**) and FZD6-Gs (**c**) complexes were shown, respectively. Color coding is annotated for each protein component. ΔBRET represents the change of bioluminescence resonance energy transfer value: ΔBRET= BRET signal (GPCR-G protein sensor) − BRET signal (only G protein sensor). Data are mean ± s.e.m. (n = 3).

Polarity describes structural, biochemical, or functional asymmetry in cells^9^. All mammalian cells experience polarity during their lifespan. There are two types of polarity: the cell intrinsic apical–basal polarity and the tissue wide polarity known as PCP^10^. FZD3/6 and their downstream Disheveled (DVL) are known to relay PCP pathways albeit the underlying mechanism is still unknown^11^. On the other hand, the PCP signaling pathway has received increasing attention attributed to its important role in orchestrating oriented cell migration and other polarized cell behaviors that are critical to many developmental processes and disorders^12^. For example, the lens of the eye maintains its polarity and transparency as it grows because it has highly ordered growth patterns such as lens fiber cell elongation, alignment and orientation, which is mediated by the PCP pathway^13, 14^. Disorganized lens fiber cell alignment causes cataract^15^.

Previous studies have shown that FZD3/6 are also associated with the growth and metastasis of various cancers, through mediating the non-canonical signaling pathways^16^. For instance, migration of breast cancer cells can be inhibited through the FZD3-dependent cAMP signaling pathway^17^; FZD6 is involved in human neuroblastomas in a β-catenin-independent manner^18^. Additionally, FZD1 also participates in cancer development and progression, such as colon cancer^19^.

Despite their important roles in cancer and other human diseases, no successful drugs have been developed for any of the ten FZDs. Previous drug discovery efforts have been focused on the anti-FZD antibody by blocking Wnt-FZD interactions but the efficacy and target selectivity is so far limited. Structural insights to elucidate the receptor activation mechanism would be valuable for probing the drug intervention hotspots and guiding the design of new tool ligands for the FZDs. Recently, Xu et al. reported the first structure of FZD and G protein complex providing structural insight into a constitutively active state of the FZD7^20^. FZD1, FZD3 and FZD6 are reported to couple to G proteins as well, but the conclusion is vague^21, 22^. Indeed, there are three gaps in our understanding of FZD signal transduction that need to be addressed: whether G protein is coupled to each FZD and which subtype; the relationship between G protein coupling and Wnt signaling pathways; and whether there is a conserved activation mechanism, such as molecular switches, that is shared by all FZDs. Here we report the cryo-electron microscopy (cryo-EM) structures for FZD1, FZD3 and FZD6, including three structures in G protein-coupled and active states, and three structures in G protein-free and inactive states. The inactive/active structure pairs showed characteristic activation mechanisms of FZDs that are distinct from other GPCRs. Our study also investigated the potential crosstalk between PCP pathway and the G protein activation of FZD6, which provides a novel understanding of the Wnt/FZD signal transduction. These comprehensive understandings of the structural and functional landscape of FZDs will provide an opportunity for developing structure-based tool ligands for FZDs’ signaling investigation.

## Results

### Three constitutively active FZDs

FZDs are classified as non-classical GPCRs with multi-domain topology and mediating signaling pathways through DVL and other transducers^6, 23^ (Extended Data Fig. 1a). Previous studies showed that FZD7 can engage Gs in a ligand-free (apo) state^20^, it remains unlocked whether FZD1, FZD3 and FZD6 can signal through heterotrimeric G proteins in the absence of an agonist. To understand the downstream G protein subtypes for FZD1, FZD3 and FZD6, we first developed bioluminescence resonance energy transfer assay (BRET2) to evaluate the constitutive activity of three FZDs’ signaling through four major Gα subunits—Gq, Gi, Gs and G12^24, 25^. This assay can detect the dissociation of heterotrimeric G proteins, which is the proximal step when the G protein signaling cascades are initiated, and it has been widely used in direct measurements of receptor-transducer coupling^26^. According to the results, we found all the three FZDs exhibited constitutive activity in G protein recruitment, in which FZD1 has a constitutive Gq activity (Fig. 1a), while both FZD3 and FZD6 have constitutive Gs activities (Fig. 1b, c). In addition, FZD6 also exhibits a relatively high constitutive activity in G12 signaling (Fig. 1c). Therefore, we assembled FZD1-Gq, FZD3-Gs and FZD6-Gs complexes for structural studies aiming to elucidate the G protein binding framework for FZDs.

### Structures of FZD-G protein complexes

To obtain stable FZD-G Protein complex for structural investigation, we assembled purified FZD proteins with mini-Gα^25^, Gβ1γ2 and the camelid antibody 35 (Nb35). Size-exclusion chromatography (SEC) and SDS-PAGE analysis reveal that purified FZD1 can form a monodispersed complex with mini-Gq (Gq); both FZD3 and FZD6 can form stable complex with mini-Gs (Gs) (Extended Data Figs. 2-4). The structures of FZD1-Gq, FZD3-Gs and FZD6-Gs complexes were finally determined at a global resolution of 3.5 Å, 3.2 Å and 3.3 Å, respectively, by cryo-EM single particle analysis (Fig. 1a-c and Extended Data Figs. 2-4). The cryo-EM maps are sufficiently clear to trace the position of the receptor, the G protein trimer and the Nb35 (Fig. 1a-c and Extended Data Figs. 2-4). No density of CRD is observed because of its flexible connection through a disordered linker with the transmembrane domain. This is consistent with our previous observation with the FZD7-Gs complex^20^. The overall structure of FZD1 consists of a hinge domain (HD, residues 275-312), the transmembrane domain (TM, residues 313-622) and an amphipathic helix (H8) (Fig. 1a and Extended Data Fig. 1b). FZD3 and FZD6 also contain the HD and TM domains but lack the H8 (Fig. 1b, c and Extended Data Fig. 1b). The overall structures of three FZD-G protein complexes are similar with the canonical GPCR-G protein complexes. Structural comparison of FZD1-Gq, FZD3-Gs and FZD6-Gs with active and inactive SMO (PDB: 6OT0^27^ and 4JKV^28^, respectively) reveals an outward movement of TM6 in these three new FZD-G protein complexes relative to inactive SMO. Such a movement of TM6 was also observed in the FZD7-Gs (PDB: 7EVW^20^) previously, suggesting that these three FZD-G protein complexes (FZD1-Gq, FZD3-Gs and FZD6-Gs) are all in active states (Extended Data Fig. 5a, b).

**Fig. 2.**
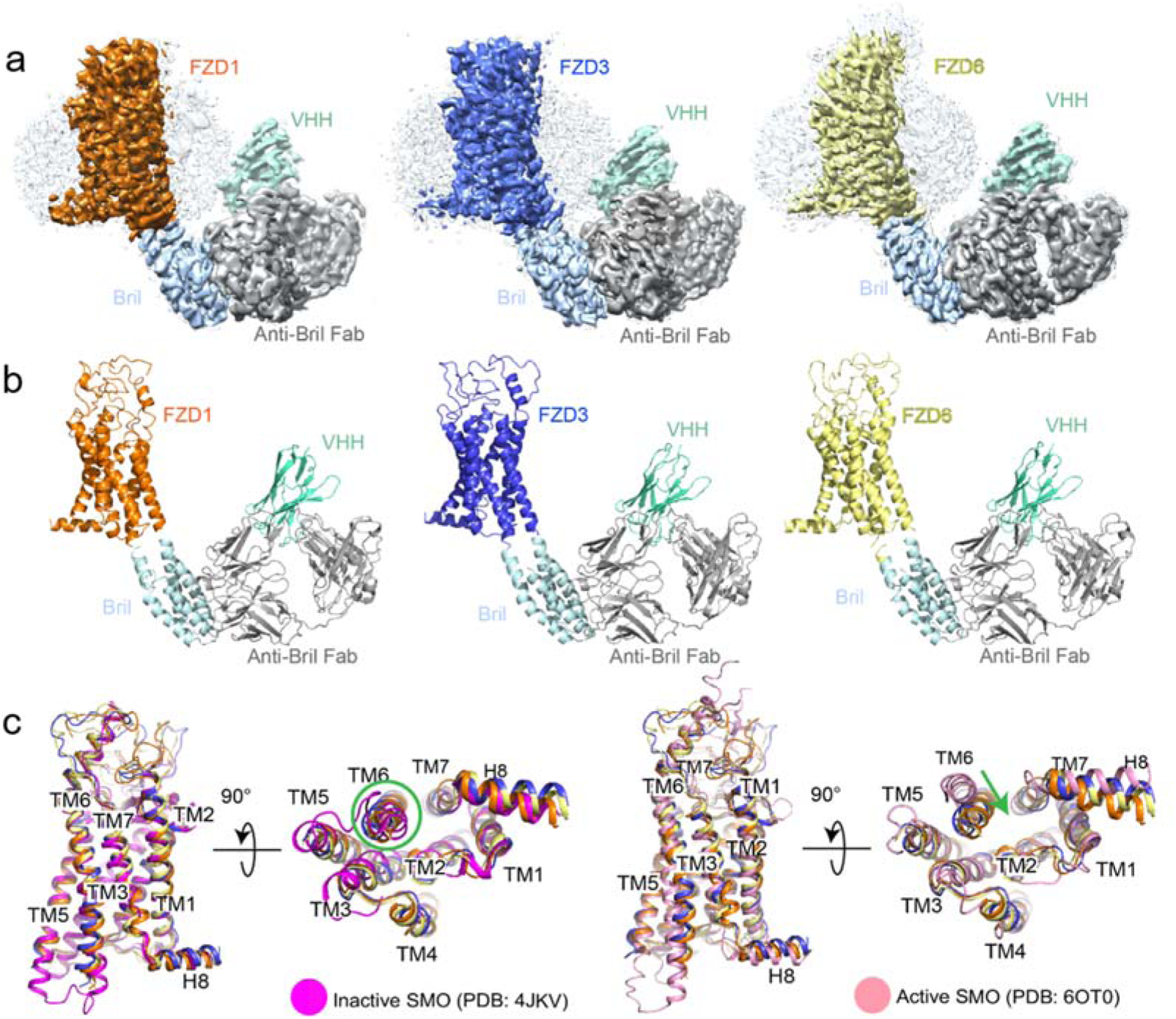
| Molecular switch for the activation of FZDs. **a, c**, Cryo-EM maps (**a**) and atomic models (**b**) of the FZD1_ICL3_BRIL-Fab-VHH (right), FZD3_ICL3_BRIL-Fab-VHH (middle) and FZD6_ICL3_BRIL-Fab-VHH (left) complexes. Color coding is annotated for each protein component. **c,** Side and intracellular views of FZD1-Fab-VHH, FZD3-Fab-VHH, FZD6-Fab-VHH with inactive (left, pink) and active (right, magenta) SMO. Transmembrane helices TM1-TM7 and helix 8 (H8) are labelled. Color coding is annotated for each protein component.

**Fig. 3.**
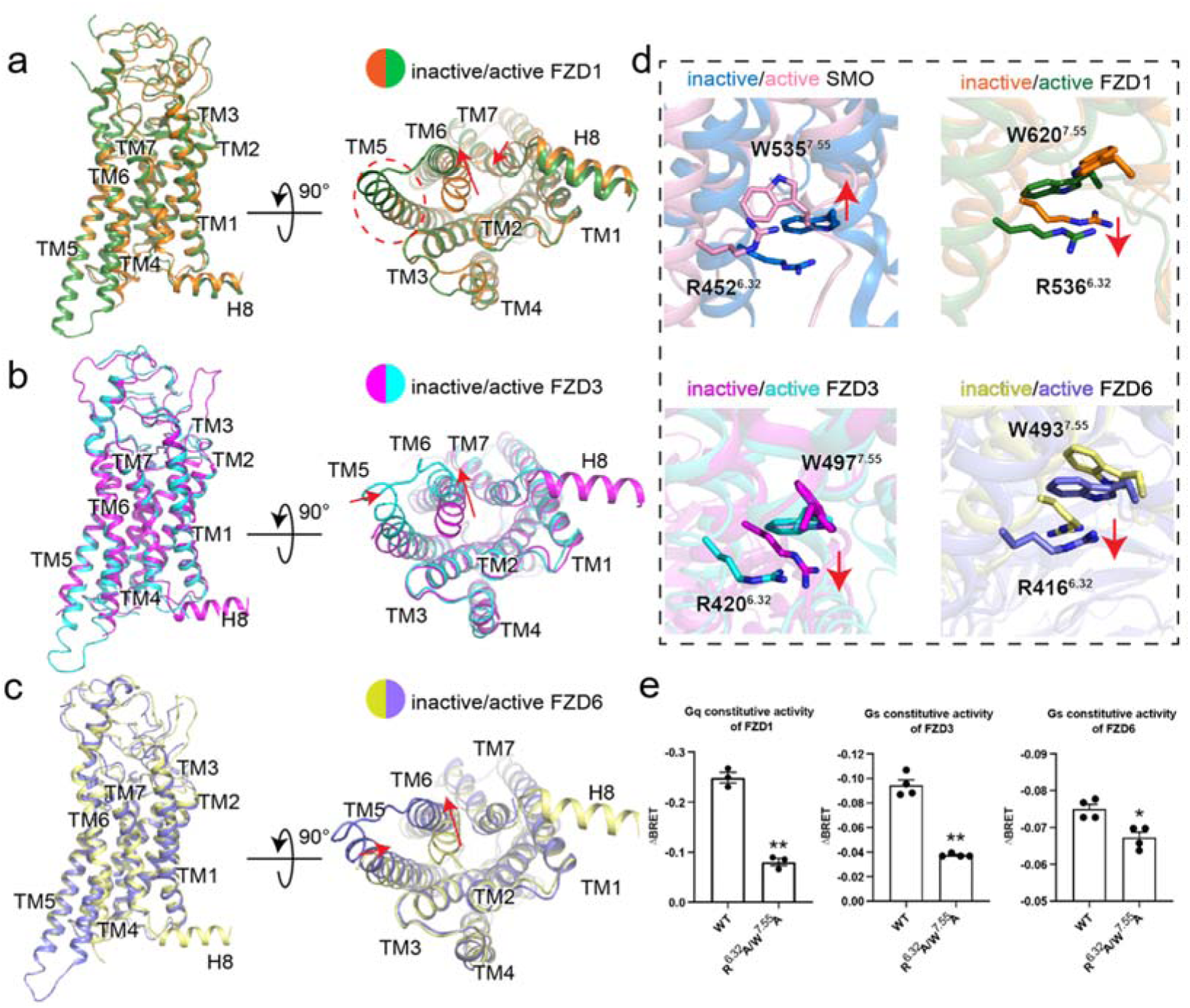
| Molecular switch for the activation of FZDs. **a-c**, Superposition of the inactive and active FZD1 (**a**), FZD3 (**b**) and FZD6 (**c**) structures. **d,** Conformational rearrangement of the R^6.32^-W^7.55^ molecular switch of SMO (inactive, 4JKV; active, 6OT0), FZD1, FZD3 and FZD6 upon G protein coupling are indicated by red arrows. Color coding is annotated for each protein component. **e,** The combined mutations of R^6.32^A and W^7.55^A reduced FZD1 (left), FZD3 (middle) and FZD6 (right) mediated G protein signaling. ΔBRET represents the change of bioluminescence resonance energy transfer value. Significance was determined by two way ANOVA with Two-stage Benjamini, Krieger, & Yekutieli FDR procedure (*P < 0.05; **P < 0.01). Data are mean ± s.e.m. (n ≥ 3).

**Fig. 4.**
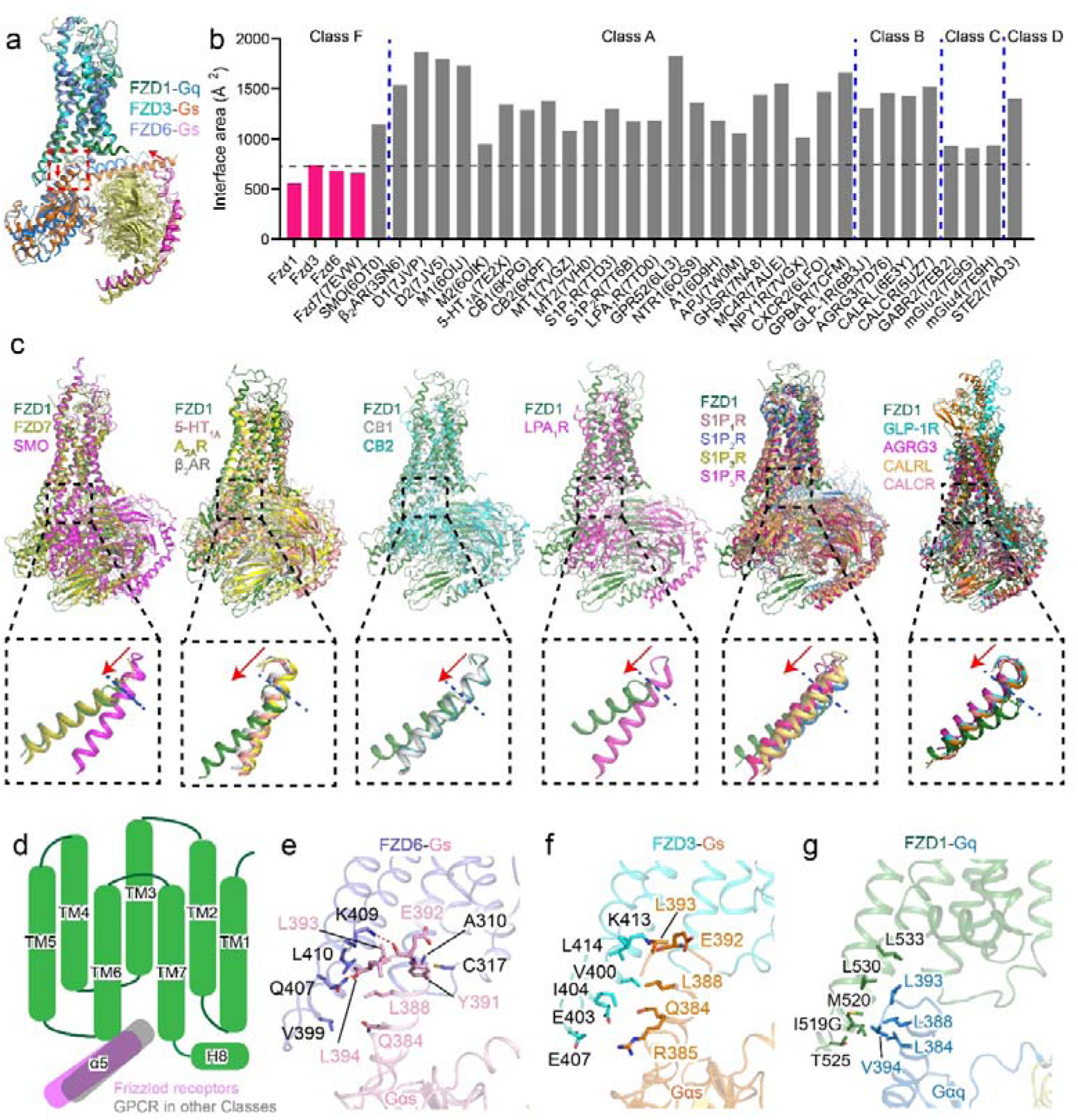
| The small interface between FZD and G proteins. **a**, Superposition of the FZD1-Gq, FZD3-Gs and FZD6-Gs structures. **b,** Calculated interface area (Å^2^) of representative GPCR-G protein complexes in five classes. **c,** Superposition of the FZD1-Gq (green) structure with representative GPCR-G protein structures. Magnified views of α5 helix of G protein are shown in respective dashed boxes with the terminal point of the α5 helix indicated by a blue dashed line. Downward movement of α5 helix in FZD1-Gq structure relative to other GPCR complexes is indicated by the red arrow. Color coding is annotated for each receptor. **d,** The schematic diagram shows the relative position of Gα protein of classical GPCRs (grey) and FZDs (pink). **e-g,** Key interactions between FZD6 and Gs (purple and pink, **e**), FZD3 and Gs (cyan and orange, **f**), FZD1 and Gq (green and blue, **g**). Key residues are shown as sticks. Color coding is annotated for each protein component.

**Fig. 5.**
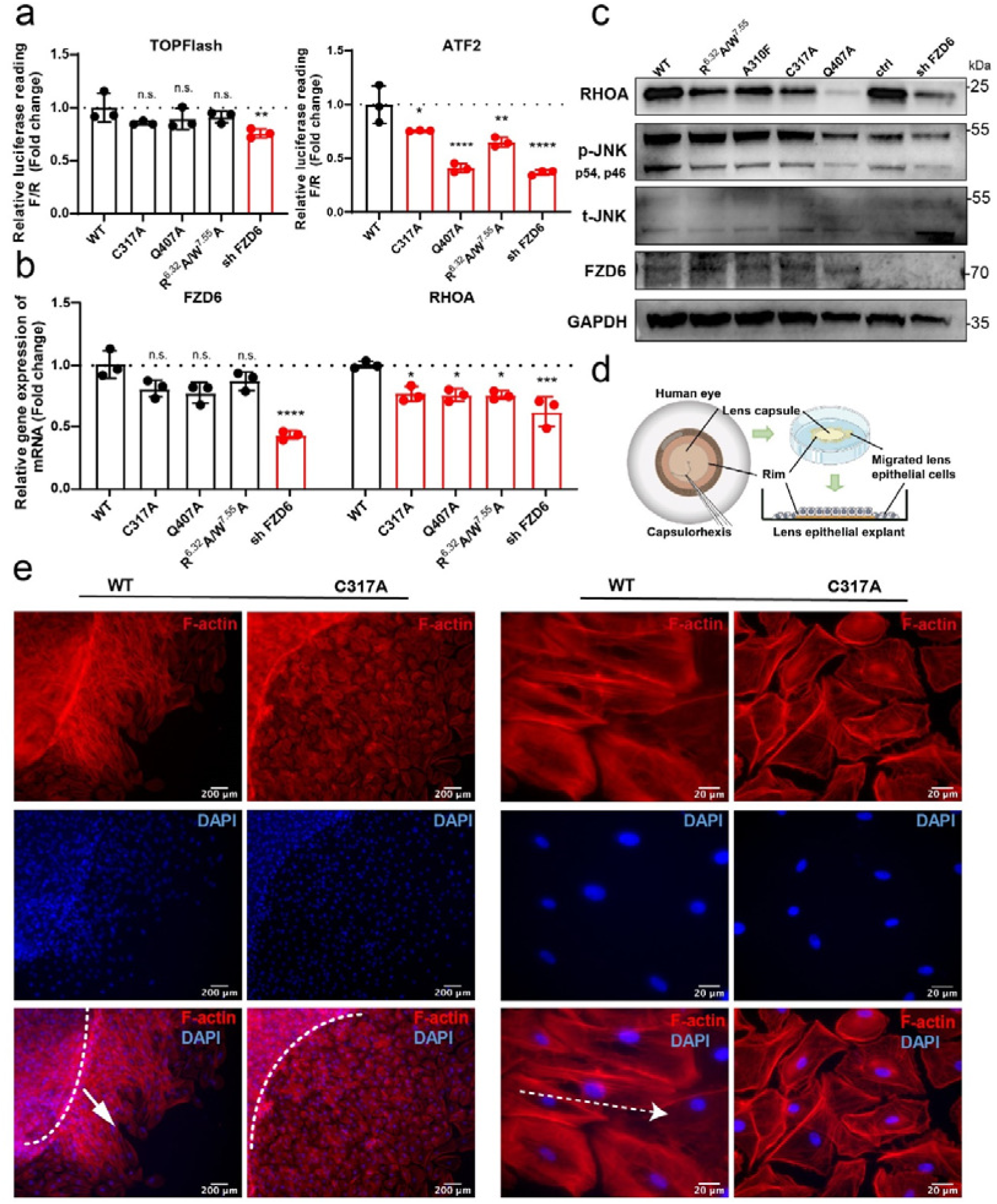
| Functional assessments of mutations on FZD6 signaling pathways in lens tissue. **a**, Relative TOPFlash and ATF2-luc reporter activity in SRA 01/04 cells transfected with control wild type (WT) FZD6, the indicated FZD6 mutants and the FZD6 shRNA; Gene reporter activities are calculated as luciferase/renilla ratios, and are normalized to WT. **b,** Quantitative RT-PCR analysis showing expression levels of FZD6 and the PCP pathway downstream gene RHOA in SRA 01/04 cells transfected with WT FZD6, the indicated FZD6 mutants and the FZD6 shRNA; Data are mean ± s.e.m. (n = 3). Significance was determined by two-way ANOVA with Two-stage Benjamini, Krieger, & Yekutieli FDR procedure (****P < 0.0001, ***P < 0.001, **P<0.01, *P<0.05, n.s. (not significant)). **c,** Western blotting assay showing protein expression levels of RHOA, phospho-JNK (p-JNK, including p46 isoform and p54 isoform), total-JNK (t-JNK) and FZD6 in SRA 01/04 cells transfected with WT FZD6, the indicated FZD6 mutants, negative control and the FZD6 shRNA. GAPDH was used as the loading control. **d,** Scheme of primary culture of human lens epithelial explant. The anterior capsule with attached epithelium was peeled off in lens surgery and cultured with the epithelium side up. **e,** Immunofluorescent staining of F-actin (red) and DAPI (blue) showed the primary cultured cells of the human lens epithelial explants treated with 200ng/mL FGF-2 and then transfected with WT FZD6 or the indicated FZD6 mutant. White dashed line shows the capsule rim. White arrow showed the direction of cell polarity. Left panel scale bar: 200 μm; right panel scale bar: 20 μm. Data are representative images of three independent experiments.

### Structures of three FZDs in the inactive state

To better understand the conformational changes during FZDs’ activation, we solved the structures of FZD1, FZD3 and FZD6 without G-protein heterotrimers. FZDs with a thermostabilized *Escherichia coli* apocytochrome *b*562RIL (BRIL)^29^ fused in ICL3, anti-BRIL Fab and anti-Fab VHH are expressed and purified separately and assembled into complexes *in vitro*^30^. Finally, the structures of anti-BRIL Fab bound FZD1, FZD3 and FZD6 (FZD1-Fab-VHH, FZD3-Fab-VHH and FZD6-Fab-VHH) were determined at a global resolution of 3.4Å, 3.3Å and 3.2Å, respectively (Fig. 2a and Extended Data Figs. 2-4). These density maps allowed us to trace the polypeptide chains and build the atomic structures of three FZDs from the hinge domain to H8, except ICL3 due to the replacement of BRIL (Fig. 2b). The CRD domain is still disordered and unmodeled in these structures, indicating that the CRD domain is flexible in both G-protein coupled and G-protein free FZDs. Through comparison of FZD1-Fab-VHH, FZD3-Fab-VHH and FZD6-Fab-VHH with active and inactive SMO as well as with inactive FZD4 (PDB: 6BD4^7^), we found that these three FZD-Fab-VHH complexes were captured in an inactive state resembling the conformation of the inactive SMO and inactive FZD4 structures (Fig. 2c and Extended Data Fig. 5c). Thus, we conclude that these three FZD-Fab-VHH complexes (FZD1-Fab-VHH, FZD3-Fab-VHH and FZD6-Fab-VHH) are all in inactive state.

### Activation mechanism of FZDs

To understand the activation mechanism of FZDs, we compared each inactive-state FZD with its respective active-state structure. Structural comparison of FZD-G protein complexes and FZD-Fab-VHH complexes reveals an outward movement in TM6 at the cytoplasmic side in the G protein-coupled relative to G protein-free states. In addition, an inward shift of TM7 in FZD1 and inward shift of TM5 in FZD3 and FZD6 were observed at the cytoplasmic side in the FZD-G protein complexes (Fig. 3a– c).

W^7.55^ and R^6.32^ have been proposed to serve as a conserved molecular switch in class F receptors including SMO^23^. For SMO, previous study suggested a weakened but preserved π-cation interaction between W^7.55^ and R^6.32^ after Gi coupling^27^. For FZDs, this molecular switch was not mentioned in the inactive FZD4 structure^7^; subsequent research proposed that this π-cation interaction in inactive FZD5 might be more extensive than that in FZD4 (Extended Data Fig. 6h)^31^; structural analysis and molecular dynamics studies of the FZD7-G protein complex suggested that the dynamic interaction between W^7.55^ and R^6.32^ might play a role in the receptor’s activation^20^. All previous conclusions are vague due to the lack of inactive/active pair for the same receptor: FZD4 and FZD5 only have inactive structures while FZD7 only has active structure. Here we compared the inactive-to-active structural transformations for FZD1/3/6 by taking advantage of their structures in both functional states reported in this study. Through structural alignment of inactive/active structures of FZD1/3/6, a consistently downward movement of W^7.55^ was observed while the three FZDs were activated (Fig. 3d). However, W^7.55^ in SMO shows an upward movement upon activation (Fig. 3d). Despite conformational changes, the strong π-cation interaction between W^7.55^ and R^6.32^ persists in the active structures for FZD1/3/6 (Extended Data Fig. 6a-c). The combined alanine mutations of W^7.55^ and R^6.32^ reduced the basal signaling activity of FZD1/3/6, implying the importance of this molecular switch for FZDs activation (Fig. 3e).

Another two conserved structural features were found by careful alignment of inactive/active pairs for each FZD: kink P^6.43^ and the W^3.43^-Y^6.40^ interaction. The outward movement of TM6 in all three FZDs begins at P^6.43^, which together with I^7.47^ and V^7.48^ forms a hydrophobic network between TM6 and TM7. When the receptor is activated, P^6.43^ kinks away from I^7.47^ and V^7.48^ and triggers the movement of TM6 at the cytoplasmic end (Extended Data Fig. 6d). Sequence alignment shows that P^6.43^, I/V/L^7.47^ and V^7.48^ are conserved in all ten FZDs Extended Data Fig. 6i), indicating a family-characteristic motif related to activation of FZDs. Mutation of P547^6.43^A in FZD1 which abolished its ability to kink decreased the Gq signaling for FZD1 (Extended Data Fig. 7a). Similarly, mutation of P431^6.43^A in FZD3 and P427^6.43^A, I485^7.47^G, V483^7.48^G in FZD6 also impaired the basal activity in Gs signaling (Extended Data Fig. 7c, e). The W^3.43^-Y^6.40^ interaction pair was also found important for FZD activation. When coupled to the G proteins, W420^3.43^ in TM3 of FZD1 moves from TM5 (inactive state) to TM6 (active state) to interact with Y544^6.40^. Such a movement of W^3.43^ was also observed in FZD3 but weakened in FZD6 (Extended Data Fig. 6f). The combined alanine mutations of W^3.43^ and Y^6.40^ reduced the basal signaling activity of FZD1 and FZD3; while no significant decrease was observed in FZD6 consistent with the structural findings (Extended Data Fig. 7a, c, e). The conformational features of these two activation motifs (kink P^6.43^ and the W^3.43^-Y^6.40^ interaction) are conserved in the inactive FZD4, inactive FZD5 and active FZD7 structures (Extended Data Fig. 6e, g) and shared by the other FZDs according to the sequence alignment (Extended Data Fig. 6i). However, the sequences of these two motifs are not conserved in SMO further indicating that SMO and FZDs may employ different activation mechanisms (Extended Data Fig. 6i).

### The small interface between FZDs and G proteins

Previous structural analysis showed a very small interface between FZD7 and Gs in the FZD7-Gs complex structure^20^, which prompted us to examine the G protein engagement in the three FZD-G protein complexes reported in this study. First, we calculated the interface area of several representative GPCR-G protein complexes and found that the four FZDs—FZD1, FZD3, FZD6, and FZD7 have the smallest G protein interface compared to receptors in other representative classes (Fig. 4a, b). It is noteworthy that the G protein interface area in SMO is about twice of FZDs (Fig. 4b), which further illustrates that SMO is different from FZDs in their activation mechanisms. Next, we compared the overall structure of FZD1-Gq complex with several representative GPCR-G protein complexes, including β_2_ adrenergic receptor (β_2_AR)^32^, adenosine A_2A_ receptor (A_2A_R)^33^, serotonin receptor (5-HT_1A_)^34^, cannabinoid receptors (CB1 and CB2)^35^, lysophosphatidic acid receptor (LPA_1_R)^36^, sphingosine 1-phosphate receptors (S1P_1_R, S1P_2_R, S1P_3_R and S1P5R)^36–39^ in class A, glucagon-like peptide-1 receptor (GLP-1R)^40^ and calcitonin gene-related peptide receptor (CALRL)^41^, the calcitonin receptor (CALCR)^42^ in class B1, and adhesion receptor AGRG3^43^ in class B2, and found that the position of the αΗ5 in FZD1 is at least one helix lower than that in other GPCRs when we align on the receptor side (Fig. 4c, d).

The overall G protein engagements of FZD3/6 are similar, which are mainly mediated by the hydrophobic interactions, polar interactions, and hydrogen bonds between αH5 of Gαs and TM5, TM6 and ICL3 of the receptor. In FZD6-Gs complex, L410^6.26^ forms major hydrophobic interactions with L388^G.H5.20^ and L393^G.H5.25^ (the generic numbering of GPCR database for Gα subunit^44^), C317^4.35^ makes hydrophobic contact with Y391^G.H5.23^, while the side chain of Q407^6.23^ and K409^6.25^ form hydrogen bonds with the carboxyl group of L394^G.H5.26^ and E392^G.H5.24^ (Fig. 4e). In FZD3-Gs complex, V400^5.72^, I404^5.76^ and L414^6.26^ form a triangular, hydrophobic core to interact with Q384^G.H5.16^, L388^G.H5.20^ and L393^G.H5.25^. The side chain of K413^6.25^ forms a hydrogen bond with the carboxyl group of E392^G.H5.24^; E403^5.75^ and E407^ICL3^ of FZD3 form polar interactions with Q384^G.H5.16^ and R385^G.H5.17^ of αH5 (Fig. 4f). Most of the residues that interact with Gs are conserved in FZDs, except C317 of FZD6, as well as E403 and E407 of FZD3 (Extended Data Fig. 8). Consistent with the structural findings, mutagenesis and cellular functional assay showed that most of the mutations at the interface reduced the Gs signaling activity of FZD3/6 (Extended Data Fig. 7d, f).

The Gq engagement of FZD1 is mainly mediated by the hydrophobic interactions. I519^5.75^, M520^5.76^, T525^ICL^^3^, L530^6.26^ and L533^6.29^ of FZD1 form hydrophobic interactions with L384^G.H5.16^, L388^G.H5.20^, L393^G.H5.25^, V394^G.H5.26^ of αH5, which results in the smallest receptor-G protein interface in the known G protein coupled FZDs structures (Fig. 4g). Mutations of I519G, M520A and L530E on the receptor side caused a reduced Gq signaling activity, indicating the importance of these hydrophobic interactions between receptor and Gq protein (Extended Data Fig. 7b).

Structural findings from the three pairs of FZD structures reported in this study shed light on a framework of FZD-G protein coupling and structural basis of the FZD activation mechanism. The small interface with a cluster of hydrophobic interactions is a representative feature for FZD1/3/6-G protein complexes, and most of the interacting residues on the interface are highly conserved among ten FZDs (Extended Data Fig. 8).

### Functional assessments of FZD6/PCP signaling in human lens tissue

Next we aimed to explore how the FZD-G protein signaling axis participates in the broader Wnt signaling pathways. Both FZD3 and FZD6 were known to signal through the PCP/tissue polarity system, but the reported FZD3 phenotype is limited to the nervous system^5^ while FZD6 expresses more broadly, including epidermal derivatives^45^, lateral ventricular^46^, neural tissues^47^, etc. The lens of the eye develops polarization structures through the highly coordinated behavior of its cells, making it an ideal model tissue for the study of the PCP pathway. The epithelial cells proliferate and their progeny migrate below the equator of the lens where they elongate and differentiate into secondary fiber cells, in which the FZD6-mediated PCP pathway plays an important role^13, 48, 49^.

We therefore performed the mutagenesis and functional assay in the human lens epithelial cell line SRA 01/04. We selected representative mutations in the FZD6-Gs interface (C317A and Q407A, separately) as well as activation switches (R^6.32^A/W^7.55^A combined mutation) for the assessment. Through TOPFlash (canonical β-catenin pathway) and ATF2-based (PCP pathway) luciferase reporter assays, we found that these residues are essential for the PCP pathway but not the canonical pathway (Fig. 5a). This implies that FZD6 may regulate the downstream PCP signaling through activating G proteins. To confirm this result, we also examined the expression levels of downstream factors of the PCP pathway, including Ras Homolog Family Member A (RHOA) and c-Jun N-terminal kinases (JNK). Quantitative RT-PCR assay revealed that all three mutations of FZD6 reduced the expression levels of RHOA, without altering the expression level of the FZD6 gene itself (Fig. 5b). Western blot analysis also indicated that these mutations impaired PCP signaling by reducing the protein expressions of RHOA and phospho-JNK, suggesting the activation of PCP/RHOA and PCP/JNK pathways were affected by FZD6-Gs complex structural integrity (Fig. 5c).

We then explored how the FZD-G protein signaling axis affects the lens fiber differentiation and polarized cell behaviors using primary culture of human lens epithelial explant. The human lens epithelial cell proliferates, migrates, differentiates and achieves proper fiber cell morphology and alignment throughout the lifetime of an individual to retain a high level of light transmission. These processes can be observed by treating human lens epithelial explants with FGF-2 to introduce fiber differentiation, followed by transfection with wild type FZD6 (WT) or FZD6-Gs interface mutation C317A (Fig. 5d). Immunofluorescent staining of F-actin indicated that the cells migrated from the rim of the capsule were highly oriented with well-organized alignment in the WT group, while those cells were less-organized and oriented diversely in the mutation group (Fig. 5e). Furthermore, the WT group showed cytoskeleton rearrangement with isotropic actin filaments, but the mutation group showed disordered cytoskeleton structure with anisotropic actin filaments (Fig. 5e).

Collectively, functional assessment of the FZD-G protein signaling axis indicated the G protein signaling mediated by FZD6 might be essential in activation of PCP signaling in regulating lens cell elongation and polarization.

## Discussion

FZDs have been long sought after as emerging targets for a range of cancers and developmental disorders, but so far no drugs have been successfully developed. Understanding the activation mechanisms of the FZDs is vital to unlock the receptor hotspots for drug interventions. However, structural and functional studies for the FZDs have been challenging for several reasons including the low expression and poor stability of the WT FZD proteins, the difficulty of Wnt purification, and whether or not G protein would bind or which subtype. Past studies on FZDs and other GPCRs mainly focus on one conformational state such as active or inactive, while the pairwise comparison of different conformations on one receptor is lacking even with the aid of alphafold. In this study, we first employed the sophisticated BRET2 assay to determine the FZD1/3/6 basal activity in coupling to their respective G protein subtypes. Next, through diligent screening of various constructs, we determined six structures of three FZDs in both active and inactive states. Together with inactive-to-active conformational comparison on each receptor, sequence conservation analysis and functional assessment, these structures reveal three FZD family-conserved molecular features (W^7.55^/R^6.32^ molecular switch, P^6.43^ kink and the W^3.43^-Y^6.40^ interaction) that may constitute the structural basis of the activation mechanism of FZDs distinct from SMO and other GPCR classes. A remarkably small G protein binding interface of FZDs may allow the FZDs to recruit G proteins at a basal level. The bound G protein subtypes for different FZDs may vary, which could be related to their different physiological functions. These observations systematically illustrate the general activation mechanism of the Class Frizzled receptors in the G protein signaling pathway, which provide some novel insight for understanding the Wnt/FZD signaling regulation and to guide drug discovery efforts targeting FZDs.

Among the ten FZDs, FZD6 and FZD3 are known to primarily mediate the non-canonical Wnt-PCP signaling pathways. In particular, FZD6 can express broadly and function on epidermal derivatives and other tissue systems suitable for investigation of cell polarity and migration in the human lens models. In this study, based on the structural observation, we first performed mutagenesis and cell-based functional assay to uncover the key residues that would govern the G protein coupling to FZD6. We then conducted systematic cell imaging and functional analysis to investigate the role of these residues on the downstream PCP pathways in the human lens epithelial cell lines and primary culture of lens epithelial explant. Intriguingly, these mutations impaired PCP signaling by affecting the downstream protein expression, and further leading to the disordered cell migration and fiber alignment. Our findings suggested that the PCP signaling pathway may rely on the activation of FZD6 by coupling to the heterotrimeric G proteins, indicating that the FZD6-G protein interface may play an important role in the alignment and polarity of lens fibers thus provide a novel hotspot for therapeutic intervention for cataract pathogenesis.

## Methods

### Cloning and expression of Frizzled receptors

The codon-optimized nucleotide sequence of human Frizzled-1 (FZD1; Uniprot ID: Q9UP38), Frizzled-3 (FZD3; Uniprot ID: Q9NPG1) and Frizzled-6 (FZD6; Uniprot ID: O60353) were synthesized by GenScript. For FZD1, the first 69 residues of FZD1 were removed. Haemagglutinin (HA) signal peptide, Flag tag, 10×His-tag and thermostabilized Escherichia coli apocytochrome b562RIL (BRIL) were added on the N terminus to enhance receptor expression and were then removed during purification with a tobacco etch virus (TEV) site. For FZD3, the first 22 residues and the last 124 residues of FZD3 and were removed. HA signal peptide, Flag tag, 10×His-tag and BRIL were added on the N terminus. For FZD6, the first 18 residues and the last 181 residues of FZD3 and were removed. HA signal peptide, Flag tag, BRIL and 10×His-tag were added on the N terminus. These three constructs were used to assemble the GPCR-G protein complex *in vitro*.

For the constructs of inactive FZD1, FZD3 and FZD6, additional modifications based on the above constructs for GPCR-G protein complex were introduced. Wild-type FZD1 (residue 70–515 and 528–637) was connected to BRIL using two short linkers derived from A_2A_ adenosine receptor (ARRQL between residue 515 and N-terminus of BRIL, and ARSTL between C-terminus of BRIL and residue 528); Wild-type FZD3 (residue 23–399 and 412–514) was connected to BRIL using two short linkers derived from A_2A_ adenosine receptor (ARRQL between residue 399 and N-terminus of BRIL, and ARSTL between C-terminus of BRIL and residue 412); Wild-type FZD6 (residue 19–395 and 408–510) was connected to BRIL using two short linkers derived from A_2A_ adenosine receptor (ARRQL between residue 395 and N-terminus of BRIL, and ARSTL between C-terminus of BRIL and residue 408).

Human FZD1, FZD3 and FZD6 were expressed in *Spodoptera frugiperda* Sf9 insect cells (Thermo Fisher) using the baculovirus method (Expression Systems), respectively. Cells were grown to a density of 2 × 10^6^ cells per ml and then infected with bocavirus (MOI=5). Cells were grown at 27 °C, collected by centrifugation 48 h after infection and cell pellets were stored at −80 °C for future use.

### Purification of Frizzled receptors

The cell pellets of FZD1, FZD3 and FZD6 protein were thawed and washed with a low-salt buffer (10 mM HEPES pH 7.5, 20 mM KCl, 10 mM MgCl_2_, protease inhibitor cocktail (Thermo Fisher)), and the supernatant was discarded by centrifugation at 35,000 x *g* for 30min. The cell pellets were followed by two washes with a high-salt buffer (10 mM HEPES pH 7.5, 1 M NaCl, 20 mM KCl, 10 mM MgCl_2_ and protease inhibitor cocktail). Before solubilization, purified cell pellets were resuspended and incubated with 2 mg ml^-1^ iodoacetamide (Sigma) at 4 °C for 30min. FZD protein was extracted from the membrane by adding HEPES, NaCl, n-dodecyl-β-D-maltoside (DDM) (Anatrace), and cholesteryl hemisuccinate (CHS, Sigma) to the membrane solution to a final concentration of 50 mM, 500 mM, 1.0% (w/v) and 0.2% (w/v), respectively, and stirred for 2 h at 4 °C. The supernatant was collected by centrifugation at 35,000 x *g* for 30 min and incubated with TALON IMAC resin (Clontech) and 20mM imidazole at 4 °C overnight.

For FZD1, the resin was centrifuged at 700 x *g* for 15min and washed with 15 column volumes of buffer I (50 mM HEPES pH 7.5, 500 mM NaCl, 5% (v/v) glycerol, 0.05% (w/v) DDM, 0.01% (w/v) CHS, 10 mM MgCl_2_ and 20 mM imidazole) and followed by 8 column volumes of wash buffer II (25 mM HEPES pH 7.5, 100 mM NaCl, 5% (v/v) glycerol, 0.03% (w/v) DDM, 0.006% (w/v) CHS and 40 mM imidazole). After the resin was washed with 8 column volumes of buffer II, the wash buffer was changed to 3 column volumes of exchange buffer (50 mM HEPES pH 7.5, 500 mM NaCl, 10% (v/v) glycerol, 0.5% (w/v) lauryl maltose neopentyl glycol (LMNG) (Anatrace), 0.1% (w/v) CHS, 10 mM MgCl_2_ and 20 mM imidazole) and incubated for 2 h at 4 °C. Then the resin was resuspended with 3 column volumes of buffer III (25 mM HEPES pH 7.5, 100 mM NaCl, 5% (v/v) glycerol, 0.03% (w/v) LMNG, 0.006% (w/v) CHS and 40 mM imidazole). TEV protease was added with a molar ratio of 1:20, and the mixture was incubated at 4 °C overnight. Next, the flow-through was collected and 3 column volumes of buffer III were added and collected. Finally, the protein solution was concentrated to ∼2 mg ml^-1^ for future use.

For FZD3 and FZD6, the resin was centrifuged at 700 x g for 15min and washed with 5 column volumes of exchange buffer (50 mM HEPES pH 7.5, 500 mM NaCl, 10% (v/v) glycerol, 0.5% (w/v) LMNG, 0.08% (w/v) CHS, 0.02% (w/v) digitonin (Biosynth), 10 mM MgCl2 and 20 mM imidazole) and incubated with 3 column volumes exchange buffer for 3h at 4 ℃. followed by 8 column volumes of wash buffer (25 mM HEPES pH 7.5, 100 mM NaCl, 10% (v/v) glycerol, 0.15% (w/v) LMNG, 0.024% (w/v) CHS, 0.006% (w/v) digitonin and 40 mM imidazole). Then the resin was eluted with 4 column volumes of elute buffer (25 mM HEPES pH 7.5, 100 mM NaCl, 10% (v/v) glycerol, 0.01% (w/v) LMNG, 0.0016% (w/v) CHS, 0.0004% (w/v) digitonin and 250 mM imidazole). Finally, the protein solution was concentrated to ∼2 mg ml^-1^ for future use.

### Cloning, expression and purification of miniGαs, miniGαq, Nb35 and VHH

miniGαs used in this paper was the same as that used in the structures of the A2AR–miniGs–Nb35^50^ GPR52-miniGs-Nb35^51^. MiniGαq was designed into a multifunctional chimera based on miniGαs (same method as miniGs/15, miniG15^52^). In brief, the specificity determinants (α5) of Gαq were transferred onto miniGαs, providing possible binding sites for Nb35 antibody to stabilize the G protein heterotrimer. The Nb35 was cloned in pET22b vector with a C-terminal 6×His tag. The anti-Fab Nb (VHH) was cloned in pET26b+ vector with an N-terminal 6×His tag. MiniGαs, miniGαq, Nb35 and VHH were expressed in *E. coli* strain BL21 (DE3) cells and purified by Ni-NTA chromatography (GenScript).

For the purification of miniGα protein^53^, the pellets from 1 L of *E. coli* culture (miniGαs or miniGαq) were resuspended in buffer (40 mM HEPES pH 7.5, 100 mM NaCl, 10% glycerol, 10 mM imidazole, 5 mM MgCl_2_, 100 μM GDP, 100 μg ml^-1^ lysozyme, 50 μg ml^-1^ DNase I, 100 μM DTT and protease inhibitor cocktail) and lysed by sonication. The supernatant was discarded by centrifugation at 35,000 x *g* for 30min and loaded onto 2 ml Ni^2+^ affinity chromatography. The column was washed with 30 ml of buffer (20 mM HEPES pH 7.5, 500 mM NaCl, 10% glycerol, 40 mM imidazole, 1 mM MgCl_2_, 50 μM GDP). The column was eluted with 6 ml buffer (20 mM HEPES pH 7.5, 100 mM NaCl, 10% glycerol, 400 mM imidazole, 1 mM MgCl_2_, 50 μM GDP). The protein solution was concentrated to a volume of 2 ml and loaded onto a Superdex200 10/600 column (GE) in buffer (10 mM HEPES pH 7.5, 100 mM NaCl, 10% glycerol, 1 mM MgCl_2_, 10 μM GDP and 1 mM TCEP). Peak fractions of miniGα protein were concentrated to 20 mg ml^-1^ for future use.

For the purification of Nb35 protein^53, 54^ and VHH^31^ protein, the pellet was resuspended from 1 L of *E. coli* culture in buffer (20 mM HEPES pH 7.5, 100 mM NaCl, 10 mM imidazole, 5 mM MgCl_2_, complete protease tablets, 50 μg ml^-1^ DNase I and 100 μg ml^-1^ lysozyme) and lysed by sonication. The supernatant was discarded by centrifugation at 35,000 x *g* for 30min and loaded onto 2 ml Ni^2+^ affinity chromatography. The column was washed with 20 ml of buffer (20 mM HEPES pH 7.5, 500 mM NaCl, 40 mM imidazole) and eluted with 6 ml buffer (20 mM HEPES pH 7.5, 100 mM NaCl, 500 mM imidazole). The protein solution was concentrated to a volume of 1 ml and loaded onto a Superdex200 10/300 column (GE) in buffer (10 mM HEPES pH 7.5, 100 mM NaCl, 10% glycerol). Peak fractions of Nb35 protein and VHH protein were concentrated to 20 mg ml^-1^ for future use, respectively.

### Cloning, expression and purification of Gβ1γ2 and anti-BRIL Fab

Both the human heterodimeric Gβ1γ2 (Gγ2 contain a C68S mutation) and anti-Bril Fab were cloned into pFastbac-Dual vector with a C-terminal 6×His tag. The human heterodimeric Gβ1γ2 was expressed in *Spodoptera frugiperda* Sf9 insect cells and the anti-Bril Fab was expressed in secreted form from *Trichuplusia ni* Hi5 insect cells using the baculovirus method (Expression Systems), respectively. Cells were grown to a density of 2 × 10^6^ cells per ml and then infected with bocavirus (MOI=5). Cells were grown at 27 °C, and cell pellets and supernatant were collected by centrifugation 48 h after infection.

For the purification of heterodimeric Gβ1γ2 protein^54^, the cell pellets from 2 L of Gβ1γ2 were thawed and resuspended to 50 ml in buffer (30 mM Tris pH 8.0, 100 mM NaCl, 5 mM MgCl_2_, 5 mM imidazole, complete protease tablets, 50 μg ml^-1^ DNase I and 100 μM DTT). Cells were broken by sonication and clarified by centrifugation (38,000 x *g* for 1 h). The supernatant was loaded onto 2 ml Ni^2+^ affinity chromatography. The column was washed with 20 ml of buffer (20 mM Tris pH 8.0, 300 mM NaCl, 30 mM imidazole, 10% glycerol and 1 mM MgCl_2_), and eluted with 6 ml buffer (20 mM Tris pH 9.0, 50 mM NaCl, 500 mM imidazole, 10% glycerol and 1 mM MgCl_2_). The elute was diluted to 60 ml in buffer (20 mM Tris pH 9.0, 50 mM NaCl, 10% glycerol, 1 mM MgCl_2_, 1 mM DTT) and loaded onto a 5 ml Q FF column (GE Healthcare) at 5 ml/min. The Q FF column was washed with 40 ml buffer (20 mM Tris pH 9.0, 50 mM NaCl, 10% glycerol, 1 mM MgCl_2_, 1 mM DTT) and eluted with a linear gradient of 50-300 mM NaCl in buffer (20 mM Tris pH 9.0, 50 mM NaCl, 10% glycerol, 1 mM MgCl_2_, 1 mM DTT). The protein solution was concentrated to a volume of 1 ml and loaded onto a Superdex200 10/300 column (GE) in buffer (10 mM HEPES pH 7.5, 100 mM NaCl, 10% glycerol, 1 mM MgCl_2_, 0.1 mM TCEP). Peak fractions of heterodimeric Gβ1γ2 protein were concentrated to 5 mg ml^-1^ for future use.

For the purification of anti-Bril Fab, the cell pellets from 1 L of anti-Bril Fab were centrifuged at 2000 x *g* for 30 min, and the 1L supernatant was loaded onto a 2 ml Ni-NTA resin. The column was washed with 15 CV of wash buffer (20 mM Tris-HCI pH 7.55, 150 mM NaCl, and 20 mM imidazole) and the protein was eluted with the same buffer supplemented with 250 mM imidazole, the protein was collected and purified over gel filtration chromatography using a Superdex 200 10/300 column (GE) equilibrated in the buffer (20 mM Tris-HCI pH 7.55, 100 mM NaCl, and 10% glycerol). Monomeric fractions were pooled, concentrated to 4 mg ml^-1^ with a 30-kDa cut-off concentrator (Millipore), and flash frozen in liquid nitrogen, then stored at −80℃ for further use.

### Complex formation for cryo-EM sample preparation

About the complex formation of Frizzled-G protein. Purified Frizzled receptor, heterodimeric Gβ1γ2, miniGαs (or miniGαq), and Nb35 were mixed in a 1:1.2:1.5:2 molar ratio followed by the addition of apyrase (1 unit), respectively. The mixture was incubated at 4 °C overnight. The FZD1-Gq (FZD3-Gs or FZD6-Gs) complex was loaded on size-exclusion chromatography (Superdex 200 10/300 GL column, GE) in SEC buffer (20 mM HEPES pH 7.5, 100 mM NaCl, 0.00075% (w/v) LMNG, 0.00025% (w/v) glyco-diosgenin (GDN, Anatrace), 0.00025% (w/v) CHS, and 100 µM DTT). Peak fractions containing FZD-G protein complex were concentrated to 2.5 mg ml^-1^ for electron microscopy studies.

About the complex formation of inactive Frizzled protein. Purified Frizzled receptor, anti-BRIL Fab and VHH were mixed in a 1:1.5:2 molar ratio, respectively. The mixture was incubated at 4 °C overnight. The FZD1-Fab-VHH (FZD3-Fab-VHH or FZD6-Fab-VHH) complex was loaded on size-exclusion chromatography (Superdex 200 10/300 GL column, GE) in SEC buffer (20 mM HEPES pH 7.5, 100 mM NaCl, 0.00075% (w/v) LMNG, 0.00025% (w/v) GDN, 0.00025% (w/v) CHS, and 100 µM DTT). Peak fractions containing complex were concentrated to 2.5 mg ml^-1^ for electron microscopy studies.

### Cryo-EM sample preparation and data collection

3 μl of the purified samples (FZD-G protein complex or FZD-Fab-VHH complex) at a concentration of around 2.5 mg ml^−1^ were applied to glow-discharged 300-mesh Au grids (Quantifoil, R1.2/1.3). Excessive sample was removed by blotting with filter paper for 3.5 s before plunge-freezing in liquid ethane using a FEI Vitrobot Mark IV at 100% humidity and 8 °C.

Five datasets (FZD3-Gs, FZD6-Gs, FZD1-Fab-VHH, FZD3-Fab-VHH and FZD6-Fab-VHH) were collected on a Titan Krios 300 kV electron microscope (Thermo Fisher Scientifics, USA) equipped with a GIF Quantum energy filter (20 eV energy slit width, Gatan Inc., USA). These five datasets were recorded by a K3 camera (Gatan) at a nominal magnification of 105,000 (calibrated pixel size: 0.832 Å/pixel) and 15 e^-^/pixel^2^/s. The movies were recorded using the super resolution counting mode by SerialEM which applied the beam image shift acquisition method with one image near the edge of each hole. A 50 µm C2 aperture was always inserted during the data collection period. The defocus ranged from –0.7 to –2.2 µm. For each movie stack, a total of 40 frames were recorded, yielding a total dose of 60 e^-^/Å^2^. FZD1-Gq datasets were collected on a Titan Krios 300 kV electron microscope equipped without an energy filter. This dataset was recorded by a K3 camera (Gatan) at a nominal magnification of 29,000 (calibrated pixel size: 0.82 Å/pixel) and a defocus range of –0.7 to –2.2 µm. For each movie stack, a total of 40 frames were recorded, yielding a total dose of 60 e^-^/Å^2^.

### Cryo-EM image processing

For FZD1-Gq complex and FZD1-Fab-VHH complex, 7,650 and 4,396 movies were recorded and processed with cryoSPARC^55^. Patch motion correction was used for beam-induced motion correction. Then, contrast transfer function (CTF) parameters for each dose-weighted micrograph were estimated by patch CTF estimation. Only images with the highest resolution of less than 4 Å were selected for further processing. A total of 6,650 and 4,242 images were selected for auto blob picking, and particles were extracted to do 2D classification. 2D class averages with diverse orientations and clear secondary features were selected as 2D templates for another round of autopicking process by cryoSPARC. A total of 935,627 and 628,701particles were selected from good 2D classification to generate the initial models. These particles and initial models were used to do 3D classification in heterogeneous refinement in cryoSPARC. 544,329 and 327,768 particles were selected for the final homogeneous refinement followed by nonuniform refinement and local refinement in cryoSPARC, resulting in density map with nominal resolution of 3.53 Å and 3.39 Å for the FZD1-Gq complex and FZD1-Fab-VHH complex (determined by gold-standard Fourier shell correlation (FSC), 0.143 criterion). Estimation of local resolution was performed in cryoSPARC.

For FZD3-Gs complex and FZD6-Gs complex, 6,801 and 11,207 movies were recorded and processed with cryoSPARC. Patch motion correction was used for beam-induced motion correction. Then, contrast transfer function (CTF) parameters for each dose-weighted micrograph were estimated by patch CTF estimation. Only images with the highest resolution of less than 4 Å were selected for further processing. A total of 6,254 and 9,565 images were selected for auto blob picking, and particles were extracted to do 2D classification. 2D class averages with diverse orientations and clear secondary features were selected as 2D templates for another round of autopicking process by cryoSPARC. A total of 1,213,683 and 1,041,782 particles were selected from good 2D classification to generate the initial models. These particles and initial models were used to do 3D classification in heterogeneous refinement in cryoSPARC. 141,786 and 530,390 particles were selected for the final homogeneous refinement followed by nonuniform refinement and local refinement in cryoSPARC, resulting in density map with nominal resolution of 3.22 Å and 3.29 Å for the FZD3-Gs complex and FZD6-Gs complex, respectively (determined by gold-standard Fourier shell correlation (FSC), 0.143 criterion). Estimation of local resolution was performed in cryoSPARC.

For FZD3-Fab-VHH complex and FZD6-Fab-VHH complex, 6,443 and 4,946 movies were recorded and processed with cryoSPARC. Patch motion correction was used for beam-induced motion correction. Then, contrast transfer function (CTF) parameters for each dose-weighted micrograph were estimated by patch CTF estimation. Only images with the highest resolution of less than 4 Å were selected for further processing. A total of 5,915 and 4,865 images were selected for auto blob picking, and particles were extracted to do 2D classification. 2D class averages with diverse orientations and clear secondary features were selected as 2D templates for another round of autopicking process by cryoSPARC. A total of 620,258 and 1,156,464 particles were selected from good 2D classification to generate the initial models. These particles and initial models were used to do 3D classification in heterogeneous refinement in cryoSPARC. 221,700 and 660,274 particles were selected for the final homogeneous refinement followed by nonuniform refinement and local refinement in cryoSPARC, resulting in density map with nominal resolution of 3.27 Å and 3.21 Å for the FZD3-Fab-VHH complex and FZD6-Fab-VHH complex, respectively (determined by gold-standard Fourier shell correlation (FSC), 0.143 criterion). Estimation of local resolution was performed in cryoSPARC.

### Cryo-EM model building and refinement

The homology models of FZD1, FZD3 and FZD6 were initially generated by Alphafold^56^. For Gs trimer and Nb35, the model 6LI3^51^ (PDB) was chosen. For BRIL, anti-BRIL Fab and VHH, the model 6WW2^31^ (PDB) was chosen. Each part of the target models was docked into the electron microscopy density map using UCSF Chimera^57^. Then, these models were used for model building and refinement against the electron density map. Subsequently, the generated model was manually adjusted in Coot^58^ followed by automatic real space refinement in real space in Phenix^59^ for several iterations. The model statistics were validated using Phenix. The final refinement statistics are provided in Extended data table 1.

### BRET2 TRUPATH assay

To measure the dissociation of Gαβγ heterotrimer directly, we applied the BRET2 assay system as reported before^24^. In brief, HEK293T cells were plated in a 6-well plate. After 2 h, cells were transiently co-transfected with plasmids encoding WT or mutated FZD together with G protein BRET probe (Gαs-RLuc8 (or Gαq-RLuc8), Gβ3, Gγ9-GFP2) using Lipofectamine 2000 reagent (Thermo Fisher). The plasmids of Gαq-RLuc8, Gβ3, Gγ9-GFP2, and FZD1 were co-transfected at a plasmid ratio of 1:1:1:2.5. The plasmids of Gαs-Rluc8, Gβ3, Gγ9-GFP2, and FZD3 (or FZD6) were co-transfected at a plasmid ratio of 1:1:1:2. 24 h after transfection, cells were distributed into a 96-well microplate (30,000–50,000 cells per well) and incubated for additional 24 h at 37 ℃. For the constitutive activity measurement, white backings (Perkin Elmer) were applied to the plate bottoms. The transfected cells were washed once with HBSS and supplemented with 100 µl of 5 µM coelenterazine 400a (Nanolight Technologies). Plates were read in EnVision plate reader (Perkin Elmer) with 410 nm (RLuc8-coelenterazine 400a) and 515 nm (GFP2) emission filters with an integration time of 1 s per well. The GFP2 emission to RLuc8 emission ratio was used to compute the BRET2 ratios.

### Transfection in SRA01/04 cell line

Lens epithelial cells were previously reported to have relatively strong constitutive expression of FZD6 and activation of WNT signalling pathways (63, 64). We adopted human lens epithelial cell line SRA 01/04 in the functional assays of downstream pathways. Cells were cultured in the Dulbecco’s Modified Eagle Medium (DMEM, Sigma-Aldrich) supplemented with 10% fetal bovine serum (FBS, #10099141, Gibco) seeded in six-well plate (for Quantitative RT-PCR and Western Blot) or 24-well plate (for Luciferase assay) overnight and were then transfected with wild type (WT) FZD6, the indicated FZD6 mutant plasmids and FZD6 shRNA. The sequence of FZD6 shRNA was: top 5’-TTTGAATTGTGCTTCAGGAAGAACTCACTTCCTGTCATGAGTTCTTCCTGAAGCACAATTATTTTT – 3’; bottom 5’-CTAGAAAAATAATTGTGCTTCAGGAAGAACTCATGACAGGAAGTGAGTTCT TCCTGAAGCACAATTC-3’.

### Luciferase assay

An ATF2-Luc reporter plasmid^60^ and TOPFlash plasmid (Beyotime) were employed to detect the activity of WNT/planar cell polarity pathway and WNT/β-catenin pathway, respectively. Cells were transfected and 300 ng of WT FZD6, mutant plasmids, or FZD6 shRNA, and then co-transfected with 300 ng of ATF2/TOP luciferase reporters and 20 ng pRL-tk. Transfections were normally carried in triplicate wells. 48 h after transfection of reporters, cells were washed twice with PBS and lysed in Passive Lysis Buffer (Promega). Luciferase activity was measured using the Dual Glo Luciferase Assay System (Promega) as instructed by the manufacturers. Gene reporter activities were calculated as luciferase/renilla ratios.

### Quantitative RT-PCR

After 48 h of transfection, cells were collected and total RNA was extracted using TRIzol reagent (Thermo Fisher), cDNA was generated using HiFiScript gDNA removal cDNA Synthesis Kit (ComWin Biotech) following the manufacturers’ instructions. Gene expression was quantified by real-time PCR with GAPDH used as an endogenous control gene. The primers used were listed here.

h-GAPDH-F ATTGCCCTCAACGACCACT

h-GAPDH-R ATGAGGTCCACCACCCTGT

h-JNK-F ACACCACAGAAATCCCTAGAAG

h-JNK-F CACAGCATCTGATAGAGAAGGT

h-RHOA-F AGGAAGATTATGATCGCCTGAG

h-RHOA-R CTAAACTATCAGGGCTGTCGAT

### Western Blot Analysis

After 48 h of transfection, SRA01/04 cell lysates were obtained using RIPA lysis buffer (Beyotime) supplemented immediately before use with 10 μl ml^-1^ phosphatase and protease inhibitors (PhosSTOPTM, Roche). The bicinchoninic acid assay (Beyotime) was used to measure total protein content to enable equal loading of protein onto 4-12% precast mini polyacrylamide gels (SurePAGE™, GenScript). Proteins were transferred onto polyvinyl difluoride (PVDF) membranes, which were then blocked with TBST containing 0.5% w/v skim milk, hybridized with primary antibody against RHOA (sc-418, Santa), phosphor-JNK (sc-293136, Santa), total-JNK (sc-7345, Santa), FZD6 (ab290728, Abcam, UK) or GAPDH (ab8245, Abcam) overnight, followed by incubation with secondary antibody conjugated with horseradish peroxidase (GE). Proteins were then detected using ECL chemiluminescent substrate (BL520A, biosharp) and visualized with a chemiluminescence gel imaging system (Peiqing).

### Lens epithelial explants collection, cell culture and treatment

All human samples were gathered from patients having lens replacement surgery at Eye & ENT Hospital of Fudan University (Shanghai, China) in accordance with the Declaration of Helsinki. By the standard step of capsulorhexis during lens surgery, the anterior capsule with attached epithelium was peeled off and collected (lens epithelium would otherwise be discarded if not for research purpose)^60, 61^. Lens epithelial explants were primarily cultured with the epithelium side up in the DMEM supplemented with 20% FBS (Gibco) in 37 °C and 5% CO_2_, with medium changed every other day. For the fibroblast growth factor 2 (FGF-2) induced lens differentiation model, explants were treated with 200 ng ml^-1^ human FGF-2 (233-10; R&D) for 48-72 hours. Explants were then transfected with WT FZD6 or the mutant plasmid.

### Immunofluorescence staining analysis

The Lens epithelial explants were fixed with 4% paraformaldehyde for 30 minutes, blocked in PBS solution containing 0.3% Triton X-100 and 3% BSA for 30 minutes, followed by incubation with TRITC Phalloidin (40734ES75, Yeasen, 1:200 for F-actin staining) and DAPI (40728ES03, Yeasen, 1:1000 for nuclei staining) at room temperature away from light for 1 hour. After washed with PBS for three times, images were obtained using a Leica DM3000 microscope system.

## Acknowledgements

This work was supported by the Ministry of Science and Technology of China 2018YFA0507000 (F.X.), the National Natural Science Foundation of China grant 32071194 (F.X.), Shanghai Science and Technology Plan 21DZ2260400 (F.X.), NSFC Outstanding Youth Foundation/National Natural Science Foundation of China grants 82122017 (X.Z.). We also thank the support from Shanghai Frontiers Science Center for Biomacromolecules and Precision Medicine at ShanghaiTech University. The cryo-EM data were collected at the Bio-EM facility at ShanghaiTech University with the assistance of Q.-Q. Sun, D.-D. Liu, L. Wang, and other staff members; We also thank the staff members of the Assay, Cell Expression, Cloning, and Purification Core Facilities of the iHuman Institute for their support.

## Author contributions

Z.Z. performed protein purification, data collection and structure determination of FZD3 and FZD6 complexes; X.L. performed protein purification, data collection, structure determination of FZD1 complexes and assisted construct design of FZD3/6_ICL3_BRIL; Z.Z. and X.L. performed structure analysis and BRET assays; L.Wei. performed luciferase assays, quantitative RT-PCR assay, Western blot and immunostaining analysis; Y.W. performed structural analysis; L.X. optimized the protein expression of FZD6 at early phase of the project; L.Wu. assisted cryo-EM data processing; X.W. cultured HEK293T cells for functional assays; X.Z. supervised the PCP signaling assays; F.X. designed, coordinated and supervised all experiments; Z.Z., X.L., L.Wei. and F.X. wrote the manuscript. All authors contributed to data interpretation and preparation of the manuscript.

## ADDITIONAL INFORMATION

Correspondence and requests for materials should be addressed to Fei Xu.

