## Supplemental files for "A framework for Frizzled-G protein coupling and implications to the Wnt-PCP signaling pathways"

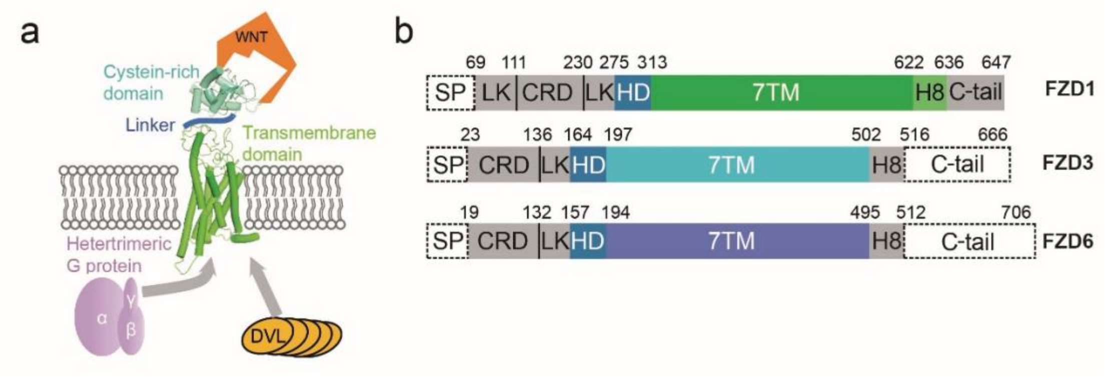


**Extended Data Fig. 1 | Schematic diagram of FZDs and schematic representation of FZD1, 3 and 6. a,** Schematic diagram of the frizzled receptors (FZDs): a flexible linker domain (blue) connects the CRD (cyan, responsible for binding to Wnt (orange)) to the transmembrane domain (green). FZDs mainly transmit signals through Dishevelled (DVL) and possibly G proteins. **b,** Schematic representation of wild-type constructs of FZD1, 3 and 6. Numbers indicate amino acid numbering in the indicated FZD subtype. The colored region stands for residues observed in respective structures. SP: signal peptide; LK: linker; HD: hinge domain. The dashed lines indicate regions not included in the constructs; the grey zones indicate regions disordered in the structures.

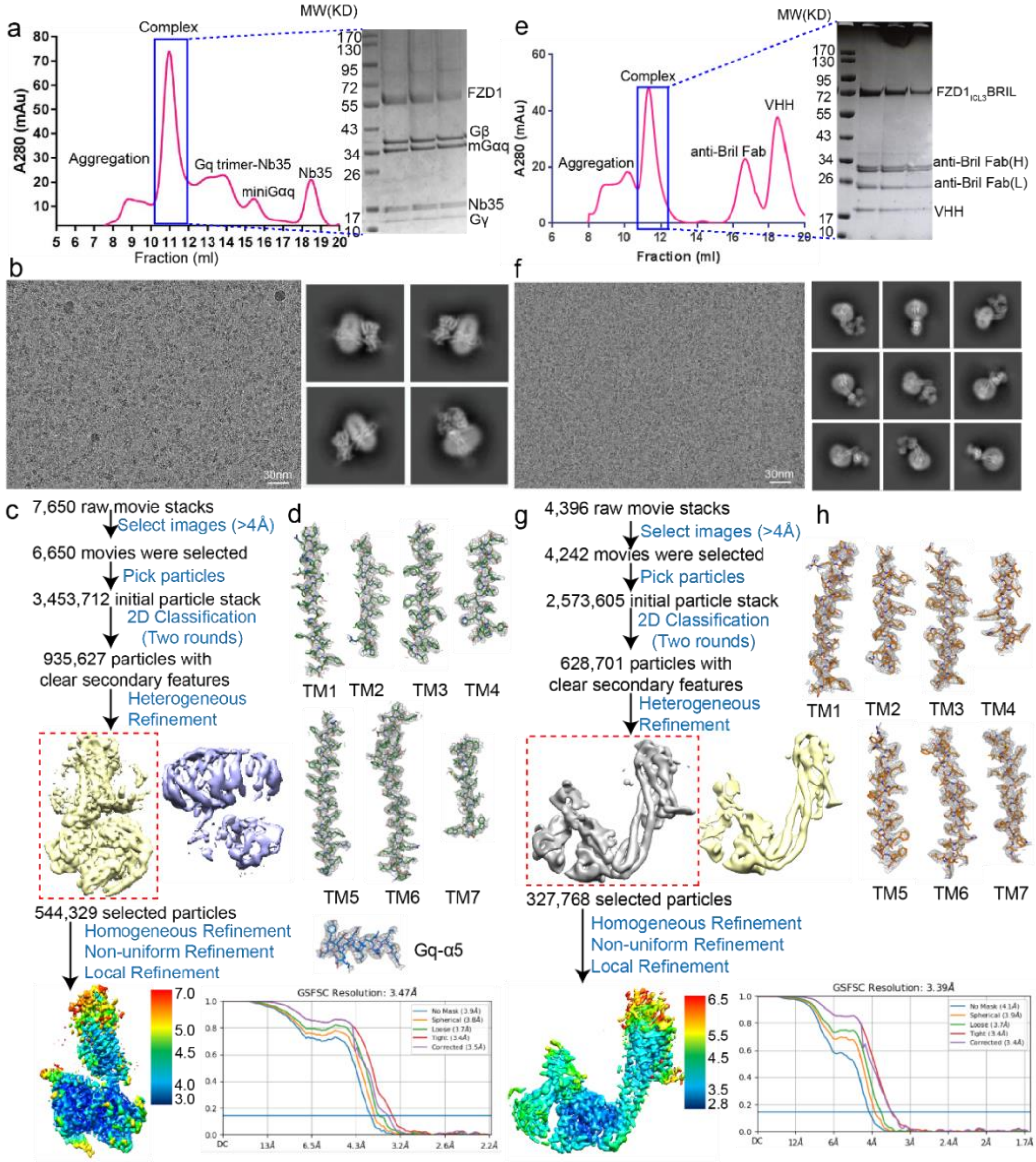


**Extended Data Fig. 2 | Cryo-EM sample preparation and data processing for FZD1 complexes. a, e,** Elution profile and gel image of FZD1-Gq (**a**) and inactive FZD1 (**e**) complexes. **b, f,** Representative cryo-EM image and representative 2D averages of the FZD1-Gq (**b**) and inactive FZD1 (**f**) complexes. **c, g,** Cryo-EM data processing workflow of FZD1-Gq (**c**) and inactive FZD1 (**g**). The data was processed by CryoSPARC and final 3D density maps are colored according to the local resolution. Gold-standard FSC curves from CryoSPARC indicate overall nominal resolutions of 3.5 Å and 3.4 Å using the FSC = 0.143 criterion for the FZD1-Gq (**c**) and inactive FZD1 (**g**) structures. **d, h**, Cryo-EM density maps and models are shown for all transmembrane helices and α5 in the Gαq protein for FZD1-Gq (**d**) and inactive FZD1 (**h**) structures, respectively.

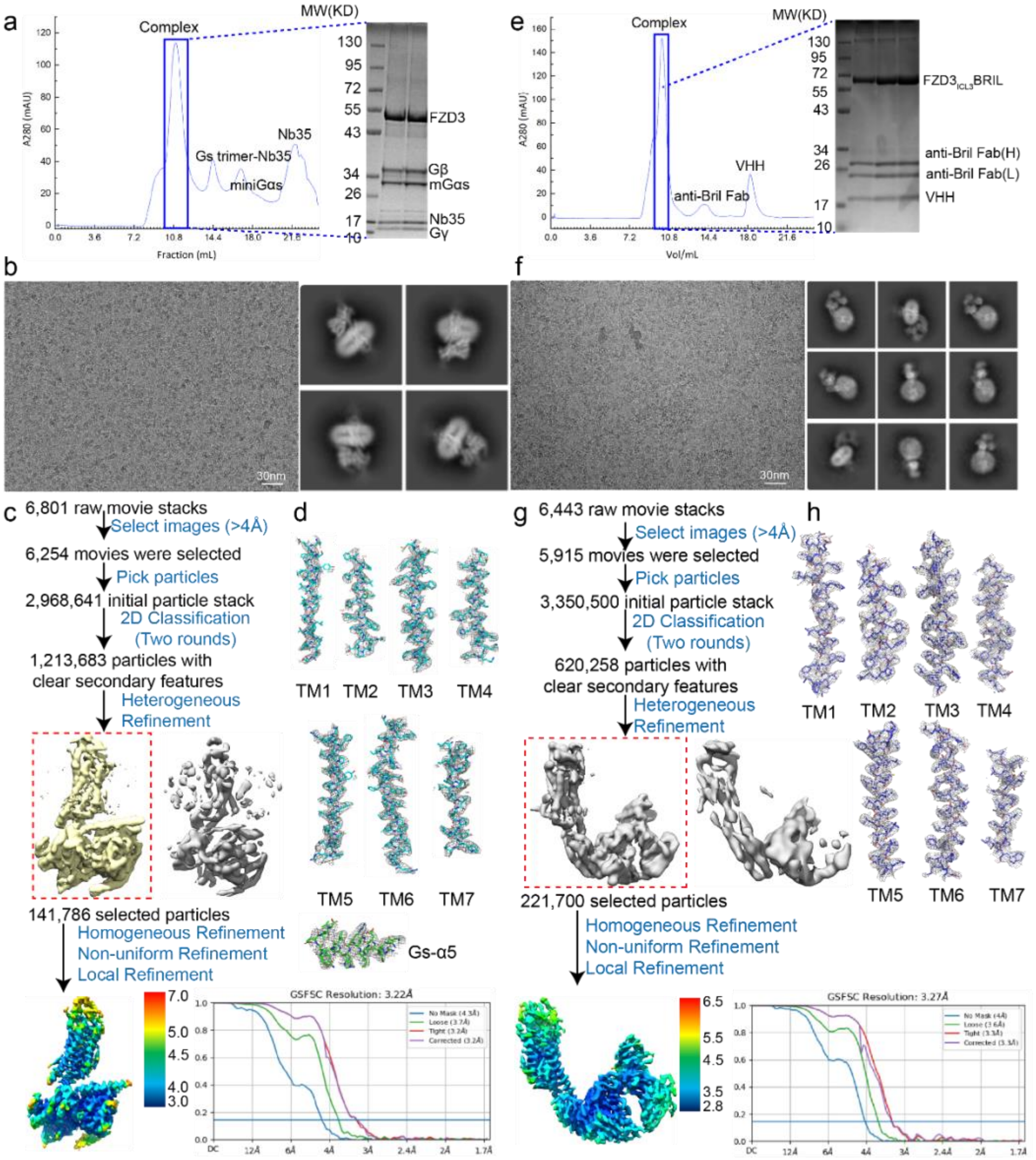


**Extended Data Fig. 3 | Cryo-EM sample preparation and data processing for FZD3 complexes. a**, **e**, Elution profile and gel image of FZD3-Gs (**a**) and inactive FZD3(**e**) complexes. **b**, **f**, Representative cryo-EM image and representative 2D averages of the FZD3-Gs (**b**) and inactive FZD1 (**f**) complexes**. c**, **g**, Cryo-EM data processing workflow of FZD3-Gs (**c**) and inactive FZD3 (**g**). The data was processed by CryoSPARC and final 3D density maps are colored according to the local resolution. Gold-standard FSC curves from CryoSPARC indicate overall nominal resolutions of 3.2 Å and 3.3 Å using the FSC = 0.143 criterion for the FZD3-Gs (**c**) and inactive FZD3 (**g**) structures. **d**, **h**, Cryo-EM density maps and models are shown for all transmembrane helices and α5 in the Gαs protein for FZD3-Gs (**d**) and inactive FZD3 (**h**) structures, respectively.

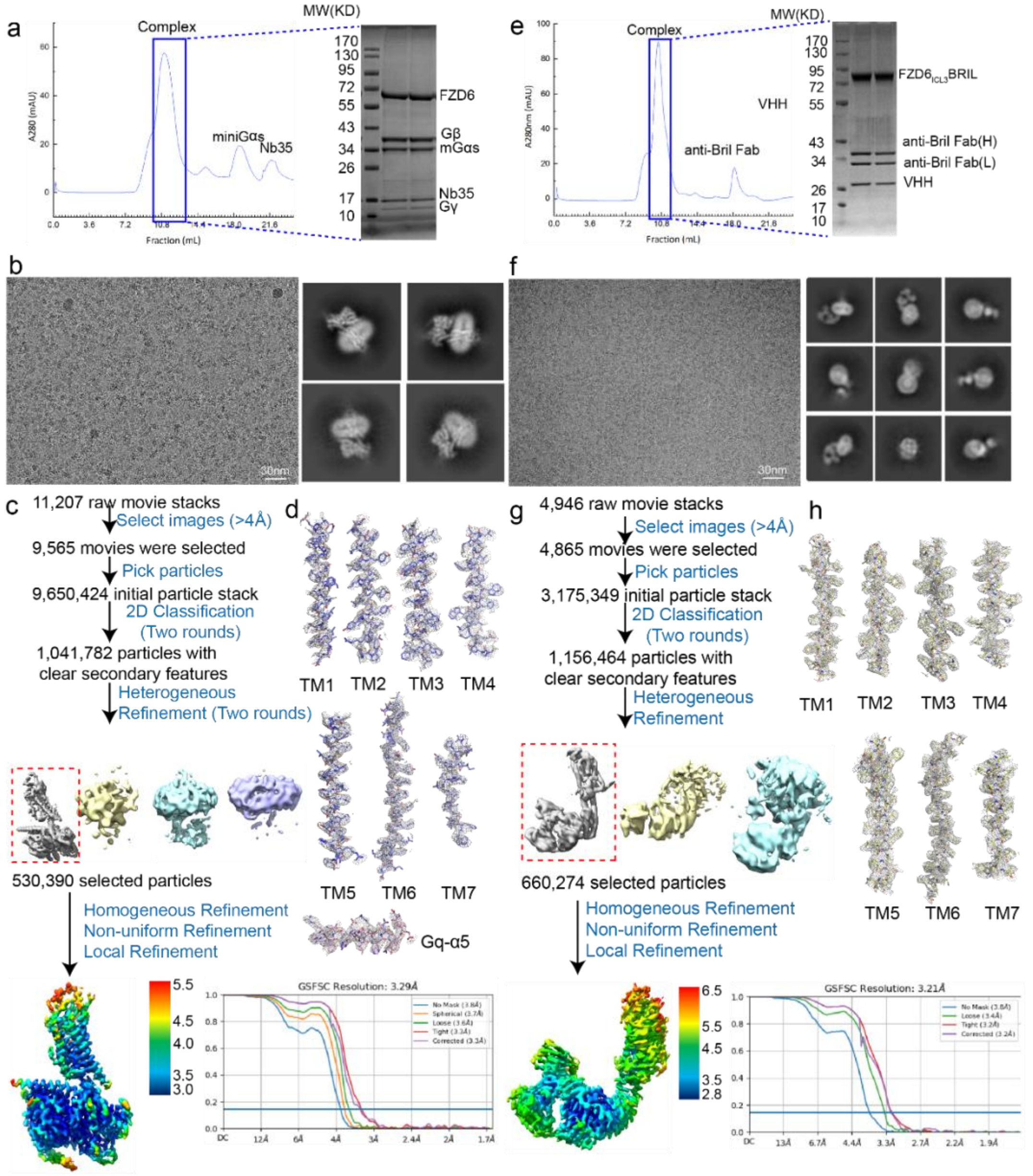


**Extended Data Fig. 4 | Cryo-EM sample preparation and data processing for FZD6 complexes. a**, **e**, Elution profile and gel image of FZD6-Gs (**a**) and inactive FZD6 (**e**) complexes. **b**, **f**, Representative cryo-EM image and representative 2D averages of the FZD6-Gs (**b**) and inactive FZD6 (**f**) complexes**. c**, **g**, Cryo-EM data processing workflow of FZD6-Gs (**c**) and inactive FZD6 (**g**). The data was processed by CryoSPARC and final 3D density maps are colored according to the local resolution. Gold-standard FSC curves from CryoSPARC indicate overall nominal resolutions of 3.3 Å and 3.2 Å using the FSC = 0.143 criterion for the FZD6-Gs (**c**) and inactive FZD6 (**g**) structures. **d**, **h**, Cryo-EM density maps and models are shown for all transmembrane helices and α5 in the Gαs protein for FZD6-Gs (**d**) and inactive FZD6 (**h**) structures, respectively.

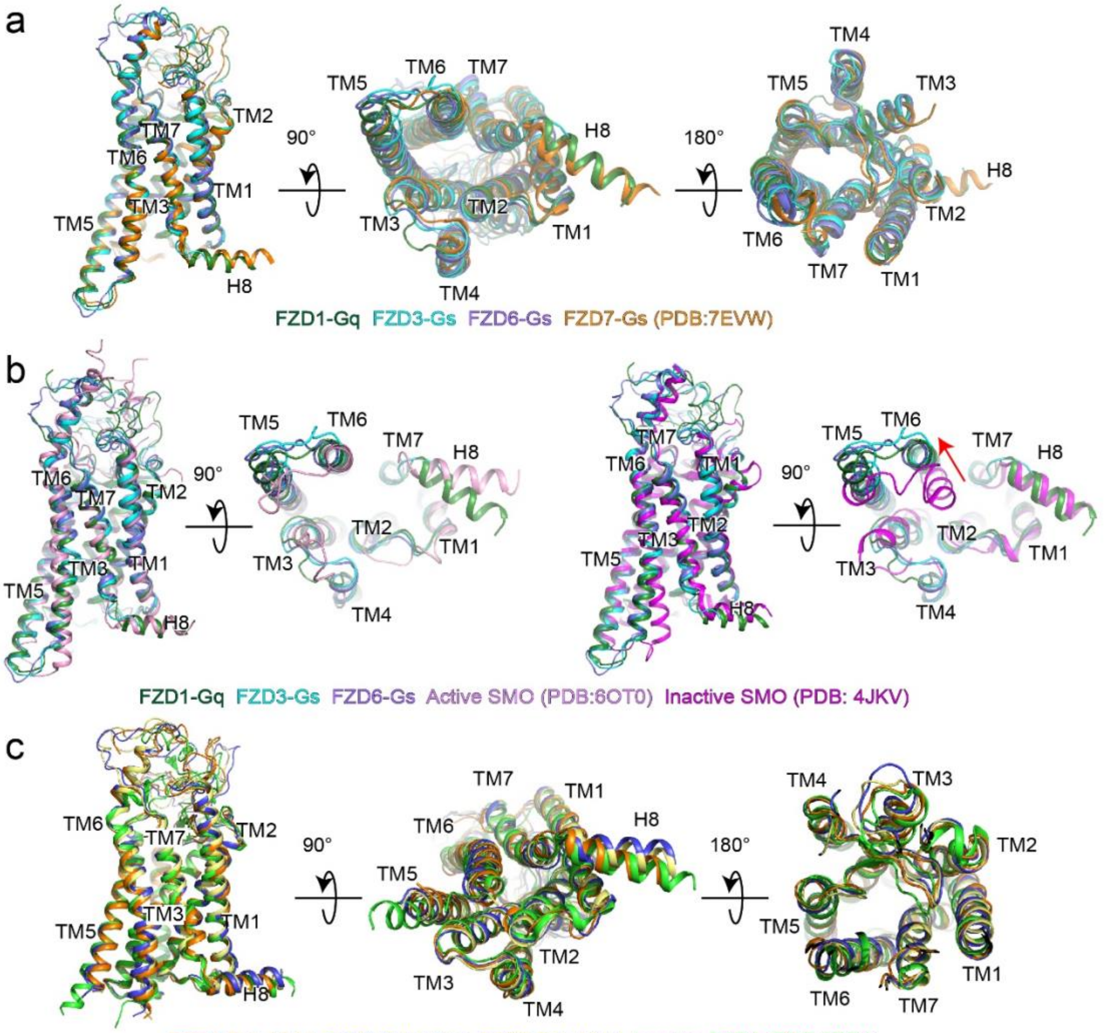


**Extended Data Fig. 5 | Structural comparison of FZD1-Gq, FZD3-Gs, FZD6-Gs complex with active and inactive smoothened receptor (SMO) and other FZD structures. a,** Side (left), intracellular (middle) and extracellular (right) views of the overlay between FZD1-Gq, FZD3-Gs, FZD6-Gs and FZD7-Gs structures. **b,** Side and intracellular views of FZD1-Gq, FZD3-Gs, FZD6-Gs with active (left, pink) and inactive (right, magenta) SMO. **c,** Side (left), intracellular (middle) and extracellular (right) views of the overlay between FZD1-Fab-VHH, FZD3-Fab-VHH, FZD6-Fab-VHH and FZD4 (green) structures. Color coding is annotated for each protein component.

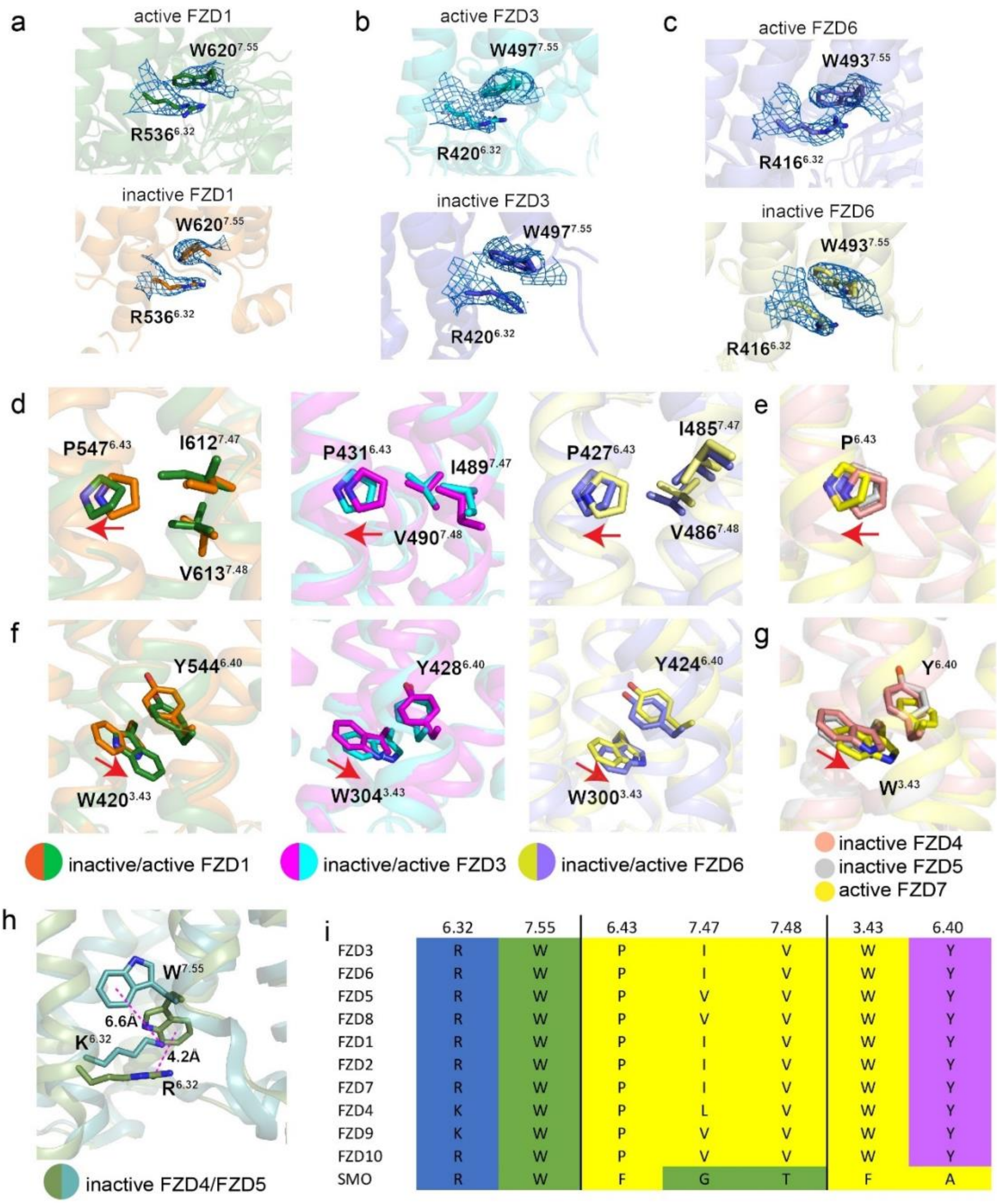


| | 6.32 | 7.55 | 6.43 | 7.47 | 7.48 | 3.43 | 6.40 |
|---|---|---|---|---|---|---|---|
| FZD3 | R | W | P | I | V | W | Y |
| FZD6 | R | W | P | I | V | W | Y |
| FZD5 | R | W | P | V | V | W | Y |
| FZD8 | R | W | P | V | V | W | Y |
| FZD1 | R | W | P | I | V | W | Y |
| FZD2 | R | W | P | I | V | W | Y |
| FZD7 | R | W | P | I | V | W | Y |
| FZD4 | K | W | P | L | V | W | Y |
| FZD9 | K | W | P | V | V | W | Y |
| FZD10 | R | W | P | V | V | W | Y |
| SMO | R | W | F | G | T | F | A |

**Extended Data Fig. 6 | Activation motifs of Frizzled receptors. a-c,** The side chains of $R^{6.32}$-$W^{7.55}$ motif in the active (above) and inactive (below) FZD1 (**a**), FZD3 (**b**) and FZD6 (**c**) structures are shown as sticks and overlaid with electron densities (blue mesh). **d,** The conformational rearrangement of residues in kink $P^{6.43}$ of FZD1 (left), FZD3 (middle) and FZD6 (right) upon G protein coupling. **e,** The conformation of residue $P^{6.43}$ in inactive FZD4 (PDB: 6BD4), inactive FZD5 (PDB: 6WW2) and active FZD7 (PDB: 7EVW) structures. **f,** The conformational rearrangement of residues in $W^{3.43}$-$Y^{6.40}$ motif of FZD1 (left), FZD3 (middle) and FZD6 (right) upon G protein coupling. **g,** The conformation of $W^{3.43}$-$Y^{6.40}$ motif in inactive FZD4, inactive

FZD5 and active FZD7 structures. **h,** The conformational comparison of R/K$^{6.32}$-W$^{7.55}$ pairs in FZD4 and FZD5. **i,** Sequence alignment of R$^{6.32}$-W$^{7.55}$, kink P$^{6.43}$ and W$^{3.43}$-Y$^{6.40}$ in all ten FZDs and SMO.

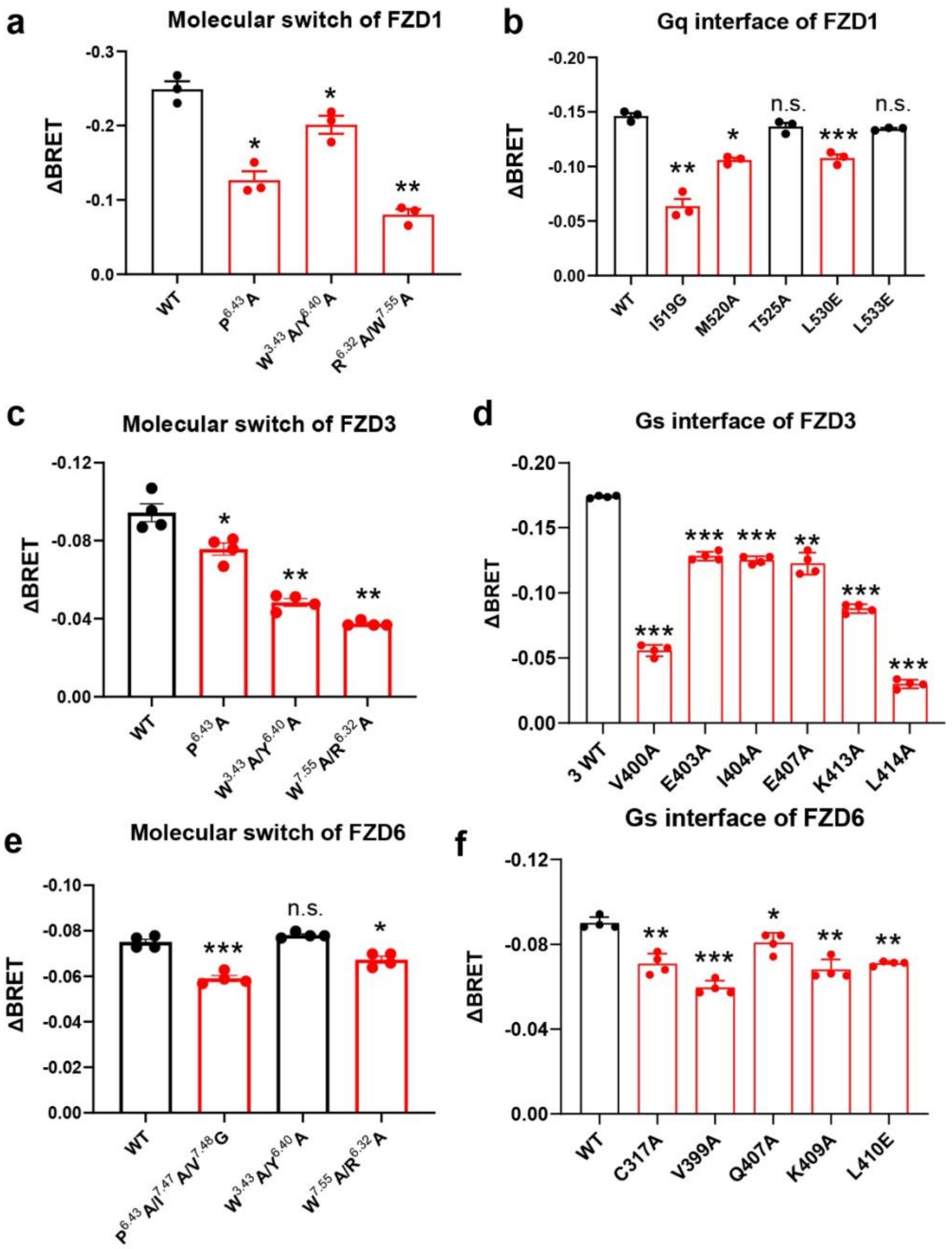


**Extended Data Fig. 7 | BRET assay of FZD mutants. a**, **b**, The effects of activation motifs (**a**) and Gq interface (**b**) mutations on FZD1's basal activity are measured by BRET assay. **c**, **d**, The effects of activation motifs (**c**) and Gs interface (**d**) mutations on FZD3's basal activity are measured by BRET assay. **e**, **f**, The effects of activation motifs (**e**) and Gs interface (**f**) mutations on FZD6's basal activity are measured by BRET assay. Significance was determined by two way ANOVA with Two-stage Benjamini, Krieger, & Yekutieli FDR procedure (***P < 0.001, **P<0.01, *P<0.05, n.s. (not significant)). Data are mean ± s.e.m. (n≥3 biologically independent experiments).

| | 12.51 | ICL2 | 4.35 | 5.72 | 5.75 | 5.76 | ICL3 | ICL3 | 6.23 | 6.25 | 6.26 | 6.29 |
|---|---|---|---|---|---|---|---|---|---|---|---|---|
| FZD3 | R | W | S | V | E | I | E | E | Q | K | L | F |
| FZD6 | R | W | C | V | V | I | D | R | Q | K | L | F |
| FZD1 | S | W | H | I | I | M | D | T | T | K | L | L |
| FZD2 | R | W | H | I | I | M | D | T | T | K | L | L |
| FZD7 | S | W | H | I | I | M | D | T | T | K | L | L |
| FZD5 | R | W | N | I | V | I | G | T | T | K | L | L |
| FZD8 | K | W | N | I | V | I | G | T | T | K | L | L |
| FZD4 | S | W | H | I | N | L | D | T | T | K | L | L |
| FZD9 | Q | W | H | I | I | M | G | T | T | K | L | L |
| FZD10 | R | W | H | I | V | M | G | E | T | K | L | L |
| SMO | R | Y | · | I | N | H | S | K | A | K | I | T |

**Extended Data Fig. 8 | Sequence alignment of ten FZDs and SMO at the G protein interface region.** Colors represent the properties of residues: blue background: basic; pink background: acidic; green background: polar; yellow background: nonpolar.

**Extended Data Table 1 | Cryo-EM data collection, refinement and validation statistics**

| | FZD1-Gq | FZD3-Gs | FZD6-Gs | FZD1-Fab-VHH | FZD3-Fab-VHH | FZD6-Fab-VHH |
|---|---|---|---|---|---|---|
| **Data collection and processing** | | | | | | |
| Magnification | 29,000 | 105,000 | 105,000 | 105,000 | 105,000 | 105,000 |
| Voltage (kV) | 300 | 300 | 300 | 300 | 300 | 300 |
| Electron exposure ($e^-$/ Å$^2$) | 60 | 60 | 60 | 60 | 60 | 60 |
| Defocus range (μm) | -0.7 to -2.2 | -0.7 to -2.2 | -0.7 to -2.2 | -0.7 to -2.2 | -0.7 to -2.2 | -0.7 to -2.2 |
| Pixel Size (Å) | 1.06 | 0.832 | 0.832 | 0.832 | 0.832 | 0.832 |
| Symmetry imposed | C1 | C1 | C1 | C1 | C1 | C1 |
| Initial particle images (no.) | 3,453,712 | 2,968,641 | 9,650,424 | 2,573,605 | 3,350,500 | 3,175,349 |
| Final particle images (no.) | 544,329 | 141,786 | 530,390 | 628,701 | 221,700 | 660,274 |
| Map resolution (Å) | 3.5 | 3.2 | 3.3 | 3.4 | 3.3 | 3.2 |
| FSC threshold | 0.143 | 0.143 | 0.143 | 0.143 | 0.143 | 0.143 |
| Map resolution range (Å) | 3.0 ~ 7.0 | 3.0 ~ 7.0 | 3.0 ~ 5.5 | 2.8~6.5 | 2.8 ~ 6.5 | 2.8-6.5 |
| **Refinement** | | | | | | |
| Map sharpening B factor (Å$^2$) | -121 | -94.6 | -133.1 | -142.6 | -111 | -81.9 |
| Model composition | | | | | | |
| Non-hydrogen atoms | 8795 | 8562 | 8649 | 7761 | 7802 | 7793 |
| Protein residues | 1110 | 1077 | 1093 | 1000 | 1001 | 1004 |
| Ligands | 0 | 0 | 0 | 0 | 0 | 0 |
| B factors (Å$^2$) | | | | | | |
| protein | 58.53 | 135.06 | 88.05 | 119.41 | 104.76 | 87.99 |
| Ligand | N/a | N/a | N/a | N/a | | |
| R.m.s. deviations | | | | | | |
| Bond lengths (Å) | 0.004 | 0.004 | 0.005 | 0.006 | 0.004 | 0.004 |
| Bond angles (°) | 0.824 | 0.867 | 0.856 | 0.960 | 0.861 | 0.868 |
| Validation | | | | | | |
| MolProbity score | 1.83 | 1.81 | 1.75 | 1.88 | 1.77 | 1.76 |
| Clash score | 7.97 | 8.27 | 8.14 | 8.66 | 6.53 | 6.86 |
| Poor rotamers (%) | 0.00 | 0.00 | 0.00 | 0.00 | 0.00 | 0.00 |
| Ramachandran plot | | | | | | |
| Favored (%) | 94.15 | 94.70 | 95.64 | 93.70 | 93.82 | 94.34 |
| Allowed (%) | 5.85 | 5.21 | 4.27 | 6.30 | 6.08 | 5.66 |
| Disallowed (%) | 0.00 | 0.00 | 0.09 | 0.00 | 0.10 | 0.00 |

**Extended Data Table 2 | Basal activity of FZD mutants, measured by BRET assays**

| | | Constructs | G-protein dissociation (ΔBRET) | P value |
|---|---|---|---|---|
| | Activation motifs | WT FZD1 | -0.2492±0.0152 | |
| | | P$^{6.43}$A | -0.1266±0.0171* | 0.0152 |
| | | W$^{3.43}$A/Y$^{6.40}$A | -0.2011±0.0172* | 0.0327 |
| | | R$^{6.32}$A/W$^{7.55}$A | -0.0802±0.0104** | 0.0015 |
| FZD1 | Gq interface | WT FZD1 | -0.1464±0.0038 | |
| | | I519G | -0.0637±0.0095** | 0.0056 |
| | | M520A | -0.1061±0.0029* | 0.0106 |
| | | T525A | -0.1367±0.0048$^{n.s.}$ | 0.1974 |
| | | L530E | -0.1079±0.0048*** | 0.0003 |
| | | L533E | -0.1346±0.0010$^{n.s.}$ | 0.0611 |
| | Activation motifs | WT FZD3 | -0.0943±0.0079 | |
| | | P$^{6.43}$A | -0.0757±0.0054* | 0.0145 |
| | | W$^{3.43}$A/Y$^{6.40}$A | -0.0484±0.0034** | 0.0060 |
| | | R$^{6.32}$A/W$^{7.55}$A | -0.0375±0.0012** | 0.0014 |
| FZD3 | Gs interface | WT FZD3 | -0.1740±0.0008 | |
| | | K413A | -0.0880±0.0029*** | P<0.0001 |
| | | E407A | -0.1225±0.0072** | 0.0014 |
| | | E403A | -0.1283±0.0029*** | 0.0001 |
| | | V400A | -0.0558±0.0037*** | P<0.0001 |
| | | I404A | -0.1253±0.0026*** | P<0.0001 |
| | | L414A | -0.0300±0.0029*** | P<0.0001 |
| | Activation motifs | WT FZD6 | -0.0750±0.0023 | |
| | | P$^{6.43}$A/I$^{7.47}$A/V$^{7.48}$G | -0.0590±0.0023*** | 0.0007 |
| | | W$^{3.43}$A/Y$^{6.40}$A | -0.0780±0.0011$^{n.s.}$ | 0.0500 |
| | | R$^{6.32}$A/W$^{7.55}$A | -0.0673±0.0026* | 0.0198 |
| FZD6 | Gs interface | WT FZD6 | -0.0900±0.0025 | |
| | | K409A | -0.0680±0.0042** | 0.0070 |
| | | Q407A | -0.0808±0.0041* | 0.0205 |
| | | V399A | -0.0595±0.0029*** | 0.0009 |
| | | L410E | -0.0710±0.0009** | 0.0010 |
| | | C317A | -0.0708±0.0042** | 0.0016 |

Data are mean ± s.e.m. from at least three independent experiments. ***P < 0.001, **P < 0.01, *P < 0.05, n.s. (not significant) by two-way ANOVA with Two-stage Benjamini, Krieger, & Yekutieli FDR procedure compared to the response of wild type.

**Extended Data Table 3 | PCP assays of FZD6 mutants**

| | Constructs | TOPFlash | P value | ATF2 | P value |
|---|---|---|---|---|---|
| WT | WT FZD6 | 1.0000±0.1350 | | 1.0000±0.1750 | |
| Control | FZD6 shRNA | 0.7568±0.0407** | 0.0083 | 0.3680±0.0221**** | <0.0001 |
| Activation motifs | R$^{6.32}$A/W$^{7.55}$A | 0.9121±0.0545 $^{\text{n.s.}}$ | 0.5093 | 0.6504±0.0434** | 0.0023 |
| Gs interface | C317 | 0.8545±0.0180 $^{\text{n.s.}}$ | 0.1303 | 0.7560±0.0050* | 0.0265 |
| | Q407A | 0.8936±0.0993 $^{\text{n.s.}}$ | 0.345 | 0.4097±0.0407**** | <0.0001 |

| | Constructs | mRNA of FZD6 | P value | mRNA of RHOA | P value |
|---|---|---|---|---|---|
| WT | WT FZD6 | 1.0040±0.1119 | | 1.0003±0.0292 | |
| Control | FZD6 shRNA | 0.4312±0.0335**** | <0.0001 | 0.6245±0.1176*** | 0.0008 |
| Activation motifs | R$^{6.32}$A/W$^{7.55}$A | 0.8731±0.0765 $^{\text{n.s.}}$ | 0.5046 | 0.7527±0.0319* | 0.034 |
| Gs interface | C317 | 0.8088±0.0721 $^{\text{n.s.}}$ | 0.1347 | 0.7622±0.0621* | 0.0442 |
| | Q407A | 0.7708±0.0812 $^{\text{n.s.}}$ | 0.0505 | 0.7545±0.0481* | 0.0357 |

Data are mean ± s.e.m. from at least three independent experiments. ****P < 0.0001, ***P <0.001, **P < 0.01, *P < 0.05, n.s. (not significant) by two-way ANOVA with Two-stage Benjamini, Krieger, & Yekutieli FDR procedure compared to the response of wild type.
